# Identification of novel oncogenic transcriptional targets of mutant p53 in Esophageal Squamous Cell Carcinoma

**DOI:** 10.1101/2023.03.12.532255

**Authors:** Sara A George, Viswakalyan Kotapalli, Pandilla Ramaswamy, Raju Kumar, Swarnalata Gowrishankar, Shantveer G Uppin, Murali D Bashyam

## Abstract

Missense mutations in the DNA binding domain of p53 are observed frequently in Esophageal Squamous Cell Carcinoma (ESCC). Recent studies have revealed the potentially oncogenic transcriptional networks regulated by mutant p53 proteins. However, majority of these studies have focused on common ‘hotspot’ p53 mutations while rarer mutations are poorly characterized. We had previously identified *SMARCD1* as an oncogenic transcriptional target of rare ‘non-hotspot’ p53 mutants detected from squamous cell carcinoma of the oral tongue (SCCOT). We now report the characterization of ‘non-hotspot’ p53 mutations from ESCC. *In-vitro* tumorigenic assays performed following ectopic-expression of ‘non-hotspot’ mutant p53 proteins caused enhancement of oncogenic properties in squamous carcinoma cell lines. Genome-wide transcript profiling of ESCC tumor samples stratified for p53 status, revealed several genes exhibiting elevated transcript levels in tumors harbouring mutant p53. Of these, *ARF6, C1QBP* and *TRIM23* were studied further due to their previously reported pro-oncogenic roles. Reverse transcription quantitative PCR (RT-qPCR) performed on RNA isolated from ESCC tumor samples revealed significant correlation of *TP53* transcript levels with those of the three target genes. Ectopic expression of wild type and several mutant p53 forms followed by RT-qPCR, Chromatin affinity-purification and Promoter-luciferase assays indicated the exclusive recruitment of p53 mutants – P190T and P278L, to the target genes leading to activation of expression. Several functional assays following knockdown of the target genes revealed a significant suppression of tumorigenicity in squamous carcinoma cell lines. Rescue experiments confirmed the specificity of the knockdown. The tumorigenic effect of the genes was confirmed in nude mice xenograft assays. This study has therefore identified novel oncogenic targets of rare ‘non-hotspot’ mutant p53 proteins relevant for ESCC besides validating the functional heterogeneity of the spectrum of tumor specific p53 mutations.

## Introduction

In India, cancer of the esophagus holds the eighth and sixth positions in terms of incidence and mortality rate among all cancers, respectively (Sung et al., 2021). Factors contributing to esophageal cancer development include genetic predisposition, age, excessive smoking and alcohol use, poor diet, obesity, Human Papillomavirus (HPV) infection and medical conditions like Barrett’s esophagus (Domper Arnal et al., 2015). *TP53* is the most frequently mutated gene in esophageal cancers (Lin et al., 2014). It encodes a transcription factor involved in the regulation of several cardinal cell processes including cell cycle, DNA damage repair, metabolism, senescence, cell death, etc. Somatic alterations in *TP53* that disrupt the tumor-suppressor activities of the protein are observed in 50% of all cancers. Nearly 90% of missense mutations, the most predominant type of alterations observed in *TP53*, occur in the sequence encoding the central DNA-binding domain (Baugh et al., 2018; Soussi & Béroud, 2001). In cancers, p53 is recurrently altered at six amino acid positions namely R175, G245, R248, R249, R273 and R282, which account for approximately a quarter of all reported mutations (referred to as ‘hotspot’ mutations) (Cerami et al., 2012; Gao et al., 2013). In addition to the inactivation of tumor suppressor function, these ‘hotspot’ mutant p53 proteins induce additional pro-oncogenic phenotypes and are thus termed ‘gain-of-function’ mutants (Muller & Vousden, 2014). Mutant p53 proteins exhibit gain-of-function properties through various means, of which transcriptional activation of oncogenes appears to be the most frequent (Kim & Deppert, 2004; Parrales et al., 2018; Strano et al., 2007). In-depth studies assessing the oncogenic effects of mutations in *TP53* are however generally restricted to the hotspot mutations while rarer mutations remain poorly characterized.

In this study, we performed genome-wide transcript analysis of ESCC tumors, including samples used in our previous study (Pandilla et al., 2013), stratified by p53 status and identified three novel oncogenic transcriptional targets of ‘non-hotspot’ mutant p53 – *ARF6, C1QBP* and *TRIM23*.

## Materials and methods

### Patient samples and Immunohistochemistry (IHC)

The study was conducted following approval from the Institutional Bioethics Committee of the Centre for DNA Fingerprinting and Diagnostics (CDFD) and the ethics committees of the respective hospitals as per the modified Helsinki Declaration (2005) (Adduri et al., 2014b; Pandilla et al., 2013). The ESCC tumors along with their corresponding matched normal samples were collected from various hospitals in Hyderabad. Details including status of p53 mutation, HPV infection, EGFR, Wnt/β-catenin status, microsatellite instability and loss of heterozygosity at different chromosomal loci were already available for all samples employed in this study (Pandilla et al., 2013). The categorization of tumors based on the nuclear-stabilisation status of p53 was performed as detailed in our previous study (Adduri et al., 2020). Samples exhibiting nuclear stabilization of the p53 protein were termed NS+ (nuclear-stabilized), a proxy for missense mutations in *TP53*, whereas those lacking stabilized p53 were termed NS-(nuclear-unstable). Formalin-Fixed Paraffin-Embedded blocks containing tumor or matched normal patient samples were cut into sections of 4μm thickness using a microtome (Leica, HistoCore RM2125 RTS) (Leica, Wetzlar, Germany). The tissue sections were processed as previously described (Bala et al., 2021). The specimens were probed with specific primary antibodies (anti-p53 (DO-1, 1:100, EMD Millipore, Calbiochem, Darmstadt, Germany), anti-ARF6 (ab77581, 1:100, Abcam, Cambridge, United Kingdom), anti-C1QBP (ab24733, 1:100, Abcam, Cambridge, United Kingdom), anti-TRIM23 (ab97291, 1:100, Abcam, Cambridge, United Kingdom)) as described earlier (Adduri et al., 2014a). The IHC stained slides were evaluated and scored by two experienced pathologists blinded to the clinical and molecular information of the samples as previously described (Bala et al., 2021). For ARF6, the slides were scored for cytoplasmic and membrane staining as 0: negative; 1: weak staining; 2: moderate staining; 3: strong staining. For C1QBP and TRIM23, the slides were scored based on the number of positively stained cells as 0: negative; 1: <25%; 2: 25-50%; 3: 50-75%; 4: >75%, as well as intensity of staining as 0: negative; 1: weak staining; 2: moderate staining; 3: strong staining. The summation of both scores yielded the final scores. Tumor DNA corresponding to exons 2-9 of *TP53* were screened for mutations by Sanger sequencing as described earlier (Pandilla et al., 2013).

### Genome-wide transcript profiling

Thirty-six ESCC tumors (19 NS- and 17 NS+) were selected for gene-expression analysis ensuring no significant variability in several parameters including patient age and gender, tumor grade and stage, status of tobacco or alcohol use, as well as molecular variables such as EGFR, FHIT, microsatellite instability, etc. Tumor samples infected with HPV were excluded from the study. Microarray based gene expression analyses were performed as described in our previous study (Kumar et al., 2018). Transcript levels of genes were also analysed using RT-qPCR to confirm that the microarray expression profiles faithfully reflected the expression status of genes. The microarray data have been deposited in the Gene Expression Omnibus (GEO) (GSE218109).

### Assessment of frequency of non-hotspot mutations in cancer databases

The frequencies of the non-hotspot and hotspot *TP53* mutations identified in ESCC tumors in the present study were measured and compared to the frequencies in three standard cancer databases – The Cancer Genome Atlas (TCGA), Catalogue of Somatic Mutations in Cancer (COSMIC) and The *TP53* database.

### Cell culture and manipulations

The p53-null cell line H1299 (a kind gift from Dr Saumitra Das, Department of Microbiology and Cell Biology, Indian Institute of Science, Bengaluru, India), AW13516 tongue cancer cell line (kindly provided by Dr Amit Dutt, Advanced Centre for Treatment, Research and Education in Cancer (ACTREC), Navi Mumbai, India), JHU-011, a p53 null laryngeal squamous cell carcinoma cell line (obtained from Dr David Sidransky, Johns Hopkins University, Baltimore, USA), HEK293T (a kind gift from Dr Rashna Bhandari, Centre for DNA Fingerprinting and Diagnostics, Hyderabad, India) were maintained at 37°C with 5% CO_2_ in DMEM supplemented with 10% FBS, 100U/ml penicillin, 100µg/ml streptomycin and 0.25µg/ml amphotericin B (Gibco, Grand Island, Nebraska, USA). Esophageal cancer cell line KYSE-410 (kindly provided by Dr Rinu Sharma, Guru Gobind Singh Indraprastha University, New Delhi, India) and NT8e, a p53-null upper aero-digestive tract cancer cell line (a kind gift from Dr Abhijit De, ACTREC, Navi Mumbai, India) were grown in RPMI with supplements. The MTT (3-(4,5-dimethylthiazol-2-yl)-2,5-diphenyltetrazolium bromide), transwell migration and colony formation assays were performed as described earlier (Adduri et al., 2020). For cell growth assay, 5×10^3^ cells were seeded per well of 6 well plates, incubated for 3 days and rinsed with 1X PBS and then trypsinized. The total number of cells per well were then calculated. All the experiments were performed in triplicate.

### Molecular genetic manipulations

For over-expression studies, wild type *TP53* cDNA was amplified from p53-GFP plasmid (Cat #11770) (Addgene, Cambridge, Massachusetts, USA) and cloned into pFN21A (Promega Corporation, Madison, WI, USA) vector using standard restriction digestion and ligation method. The missense mutant constructs were generated through PCR-based site directed mutagenesis (SDM). For immunofluorescence studies, pEGFP-N1, wild-type p53-EGFP plasmid, and p53-EGFP mutant constructs generated through SDM were employed. Promoter fragments corresponding to wild type and mutant p53 target genes were amplified by standard PCR using DNA from healthy donors and subsequently cloned into the pGL4.2 basic vector (Promega Corporation, Madison, USA) for promoter-luciferase assays. p53-pFN21A expression constructs were transfected into H1299, AW13516 and KYSE-410 cells using Lipofectamine-2000 (Invitrogen, Massachusetts, USA). For rescue experiments, the coding sequences of ARF6, C1QBP and TRIM23 were amplified from pLenti6.3/V5 DEST vector (Cat. #HsCD00939228 (ARF6), #HsCD00943732 (C1QBP), #HsCD00941058 (TRIM23)) (DNASU, Arizona State University, Arizona, USA) and cloned into the pEGFP-N1 vector (Cat. #6085-1) by restriction digestion.

The short-hairpin loop RNAs (shRNAs) targeting *ARF6, C1QBP* and *TRIM23* were selected from the RNAi consortium shRNA library hosted by the Broad Institute (Clone ID: *ARF6*: TRCN0000294069, TRCN0000305862, TRCN0000379678, TRCN0000379907 and TRCN0000305860, *C1QBP*: TRCN0000057106, TRCN0000057104, TRCN0000370165, TRCN0000057107 and TRCN0000370164, and *TRIM23*: TRCN0000233071, TRCN0000233067, TRCN0000233068, TRCN0000233070 and TRCN0000233069). shRNA stable cell lines were generated as described earlier (Adduri et al., 2020).

For rescue experiments, *ARF6, C1QBP* and *TRIM23* expression constructs were generated in pEGFP-N1 vector using standard cloning procedure. Further, synonymous mutations in the region encoding three consecutive amino acids corresponding to the shRNA seed region in each of the three constructs were generated (to prevent/ reduce degradation of the target gene transcripts under the influence of shRNA) through site-directed mutagenesis (Bala et al., 2021).

### Assessment of sub-cellular localization of p53-Halo proteins

H1299 cells were transfected with pEGFP vector or p53-wild type/ mutant-pEGFP constructs and incubated for 36 hours. The slides were washed once with 1X PBS, fixed for 10 minutes with 4% paraformaldehyde and subsequently washed thrice with 1XPBS. 5% Bovine serum albumin was added for blocking and then the slides were incubated for an hour on a rocker at room temperature. The slides were washed thrice with 1X PBS, then mounted with Vectashield Antifade Mounting Medium with DAPI (4′,6-diamidino-2-phenylindole) (Vector Laboratories, Burlingame, California, USA) and sealed, followed by incubation in the dark for 30 minutes. The slides were visualized under 65X magnification in the Leica SP8 (Leica Microsystems, Wetzlar, Germany) confocal microscope. The localization of each tagged-protein was assessed with DAPI as a reference for the nucleus.

### Sodium Dodecyl Sulphate-Polyacrylamide Gel Electrophoresis (SDS PAGE) and Western blotting

In order to evaluate the relative protein levels of the ectopically-expressed genes in cell lines, SDS-PAGE and western blotting were performed as described earlier (Adduri et al., 2020) using antibodies against p53, HaloTag (Promega Corporation, Madison, USA), GFP (Life Technologies Private Limited, Bangalore, India), α-tubulin (Sigma-Aldrich, St Louis, Missouri, USA), GAPDH (Invitrogen, Carlsbad, USA), ARF6, C1QBP and TRIM23.

### Dual-luciferase reporter and Chromatin Affinity Purification (ChIP) assays

Luciferase reporter assays were performed similar to our previous study (Kumar et al., 2018). ChAP assays were performed following transfection with various p53 expression constructs using the Halo-ChIP system (Promega Corporation) according to the manufacturer’s protocol as described previously (Adduri et al., 2020). The promoter region evaluated for each gene in the ChAP experiment included the region analysed in the promoter-luciferase assays. Promoter enrichment after immunoprecipitation was measured by RT-qPCR using a standard protocol as enumerated in our previous studies (Adduri et al., 2020; Kumar et al., 2018).

### *Ex-vivo* tumorigenesis assays

AW13516 cell lines stably expressing either non-targeting scrambled shRNA (#1864) (Addgene, Cambridge, Massachusetts, USA) or shRNAs targeting *C1QBP* or *TRIM23* were employed to study the effect of knockdown on the tumorigenic potential of cancer cells. 5×10^6^ cells were mixed with 100μl of 1XPBS and 100μl of Matrigel (Corning, New York, USA) and injected into the flanks of 5-8 weeks old *FOXN1-/-* nude mice (seven and nine mice per batch for *C1QBP* and *TRIM23*, respectively) as previously reported (Animireddy et al., 2021; Bala et al., 2021). The dimensions of the tumors were measured at weekly intervals for a period of eight weeks post-injection. The mice were then necropsied and the tumor sizes and weights were measured.

## Results

### Genome-wide transcriptome analysis of ESCC tumors reveals novel transcriptional targets of mutant *TP53*

In order to identify the transcriptional targets of ESCC-specific p53 mutants, microarray-based transcript profiling was performed on a cohort of thirty-six (nineteen p53 nuclear negative (NS-) and seventeen p53 nuclear positive (NS+)) ESCC samples (Table S1) as previously described (Adduri et al., 2020). Following computational analysis of the microarray data, we identified several gene-sets (Adduri et al., 2020) differentially enriched in p53 NS+ tumors. The top forty differentially enriched gene-sets (p<0.0007) represented several pro-tumorigenic processes including p53 regulated hypoxia pathway, Wnt signalling, Myc, ERBB/EGFR signalling, etc. (Figure S1A).

Application of Significance Analysis of Microarrays (SAM) on all 1024 genes that constituted the 40 gene-sets, revealed seven genes that were significantly upregulated in NS+ samples (Fig S1B-C). Interestingly, the transcript levels of *TP53* itself were significantly up-regulated in NS+ samples (Figure S1B), validating our previous results obtained for squamous cell carcinoma of the oral tongue (SCCOT) (Adduri et al., 2020). Among the genes that appeared to be differentially upregulated in NS+ samples, three namely *ARF6, C1QBP* and *TRIM23*, were chosen for further studies based on their purported role in cancer. Transcript levels of the three genes were significantly higher in NS+ (compared to NS-) samples as confirmed through RT-qPCR (Figure 1A and Table S2A). Interestingly, the expression levels of the three genes also significantly correlated with the transcript levels of *TP53* (Figure 1B and Table S2B).

**Figure 1.**
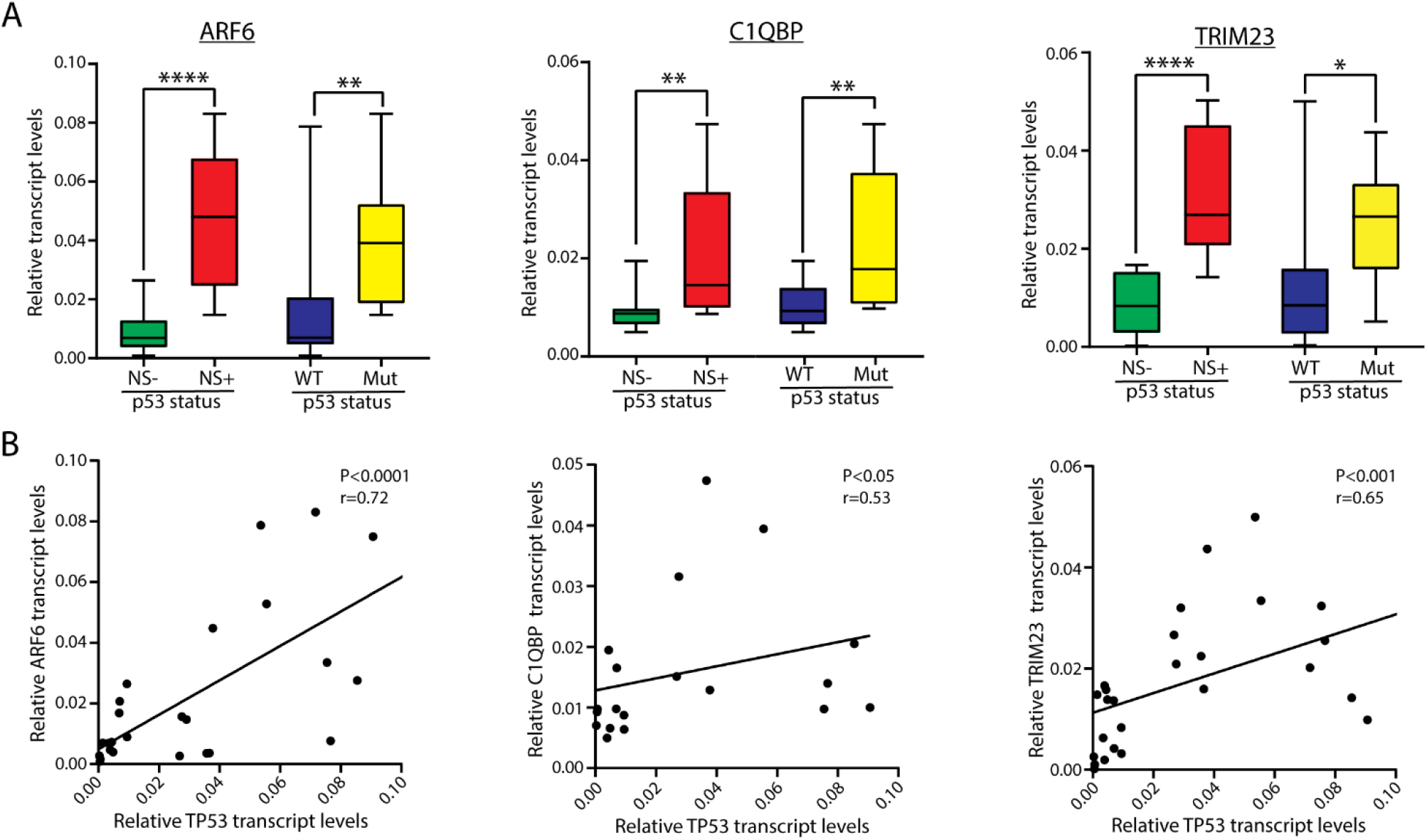
Validation of novel transcriptional targets of non-hotspot mutant p53 forms. Panel A shows relative transcript levels of the three genes (indicated) in ESCC tumors stratified for p53 nuclear stabilization or mutation status. Panel B shows correlation of transcript levels of the three genes with that of *TP53* in ESCC tumor samples. NS, nuclear stabilization. *, P<0.05; **, P<0.01; ****, P<0.0001; (unpaired student’s t-test).

Next we identified eleven different *TP53* mutations in the ESCC NS+ samples subjected to gene expression analysis (Table S1). No *TP53* mutation was detected in the ESCC NS-samples, as expected. Of note, several of these mutations were comparatively rare (Table S3) with significantly lower frequencies in cancer mutation databases. We confirmed the differential expression of the three putative target genes in p53 mutant and wild type samples (Figure1A and Table S2C). Of the various p53 mutants detected in the ESCC tumor samples, R158P, P190T, Y236C and P278L (termed ‘non-hotspot’ mutations) were further assessed for their oncogenic potential in the p53 null H1299 squamous carcinoma cell line; four highly oncogenic hotspot mutant proteins namely R175H, R248W (also detected in the ESCC tumor samples; Table S1) and R273H were employed as controls. Of note, all p53 mutant forms predominantly exhibited a nuclear localization (Figure S2). As expected, the hotspot p53 mutants supported significantly elevated cell viability/ proliferation as well as migration potential whereas, the wild type p53 supported significantly lower activities in the same assays (Figures 2A-B) (Manoochehri et al., 2014). More importantly, the non-hotspot mutants P190T and P278L consistently supported significantly high cell viability/proliferation and migratory activity (Figure 2A-B).

**Figure 2.**
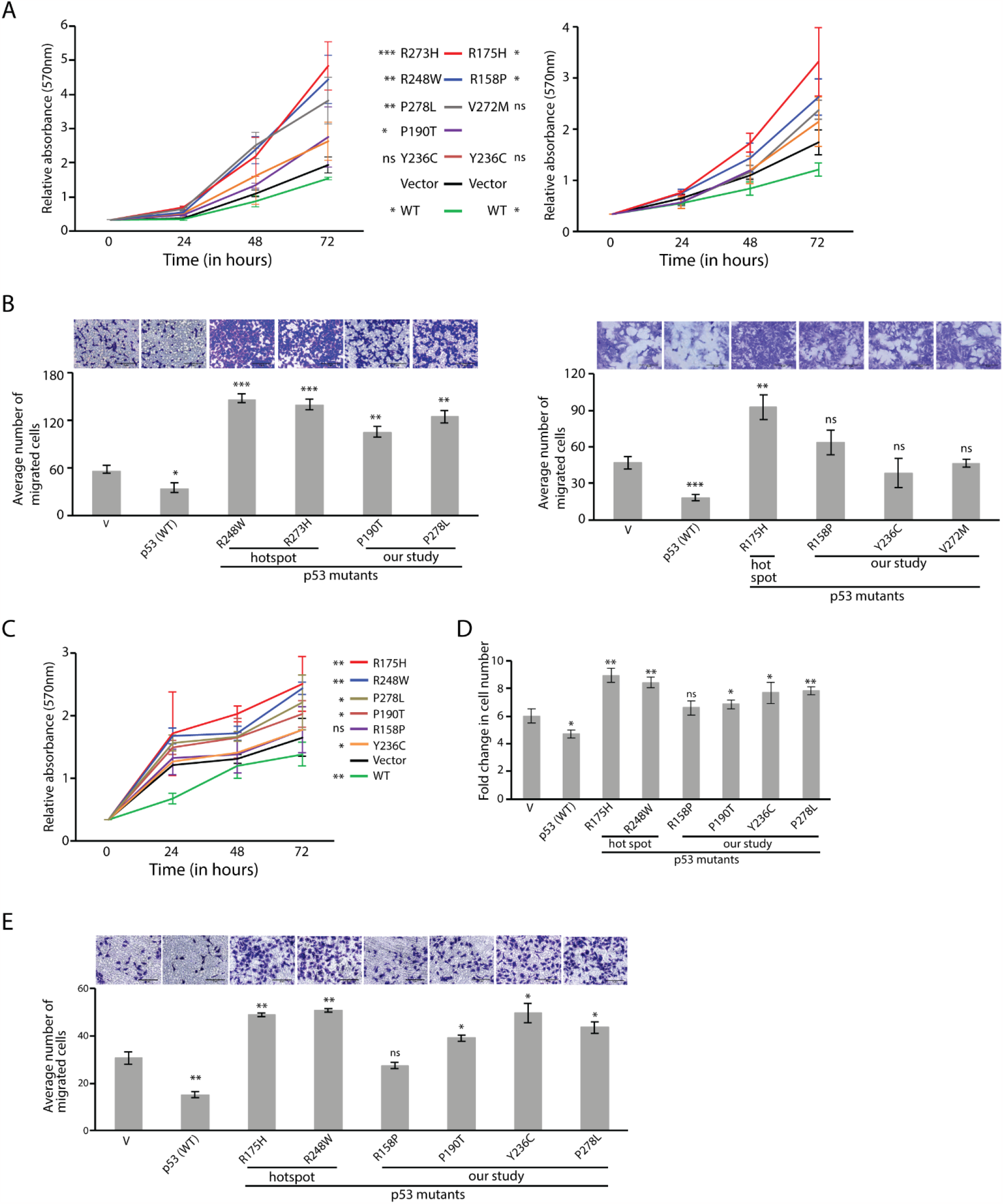
Evaluation of oncogenic potential of ‘non-hotspot’ mutant p53 proteins. Results for MTT (panels A and C), migration (panels B and E) and growth (panel D) assays performed in H1299 (panels A and B) and KYSE-410 (panels C-E) cells are shown. V, expression vector. Statistical significance is calculated on the difference between each p53 construct and vector based on at least three independent experiments; *, P<0.05; **, P<0.01; ***, P<0.001; ns, not significant (unpaired student’s t-test).

Based on the above results we proceeded to scrutinize the activity of the P190T and P278L non-hotspot mutants in additional squamous cancer cell lines namely NT8e and JHU-011 (both p53 null Head and Neck Squamous Cell Carcinoma cell lines), to validate the results obtained in H1299. Indeed, P190T and P278L mutants supported significant induction of tumorigenic activity in both cell lines, similar to the hotspot mutants (Figures S3 and S4). Finally, we evaluated the tumorigenic potential of P190T and P278L in comparison to other non-hotspot and hotspot mutations in the esophageal squamous cancer cell line KYSE-410. Both P190T and P278L induced tumorigenic activity as evidenced from three different assays namely MTT, cell growth and cell migration (Figure 2C-E).

Given that *TP53* hotspot mutations exert their pro-tumorigenic function via non-canonical transcriptional activation of oncogenes (Pfister & Prives, 2017); we endeavoured to determine whether the oncogenic potential exhibited by non-hotspot mutations P190T and P278L could be via transcriptional activation of *ARF6, C1QBP* and *TRIM23*. To this end, we first performed ectopic expression of wild type and several mutant (R158P, R175H, P190T, Y236C, R248W, P278L and R282W) p53 forms and evaluated transcriptional activation of *ARF6, C1QBP* and *TRIM23* in H1299, AW13516 as well as in KYSE-410 cells. *CDKN1A* and *MVK*, known transcriptional targets of wild type and hotspot mutant p53 forms, respectively, were evaluated as controls. RT-qPCR analyses revealed that only P190T and P278L consistently activated the expression of the target genes (Figure 3A-C). As expected, *CDKN1A* and *MVK* were exclusively activated by wild type and hotspot mutant p53 forms, respectively. We also evaluated the ability of varied p53 forms to activate the promoters of the three target genes in H1299 and KYSE-410 cells. P190T and P278L (but not the hotspot or wild type forms) were able to increase the activity of *ARF6, C1QBP* and *TRIM23* promoters (Figures 3D-E). As expected, wild-type p53 exclusively induced the expression of the luciferase gene linked to the *ATF3* promoter, while the *MVK* promoter-linked luciferase expression was regulated exclusively by the hotspot mutant R248W (Figures 3D-E). Finally, ChAP assay was performed in H1299 cells to assess whether the regulation of the target genes occurred due to the recruitment of p53 to the respective promoters. Indeed, P190T and P278L were recruited to the three target gene promoters (Figure 3F); this enrichment was promoter specific as a similar experiment targeting the gene body region of the three genes revealed a markedly lower enrichment, if any (Figure S5).

**Figure 3.**
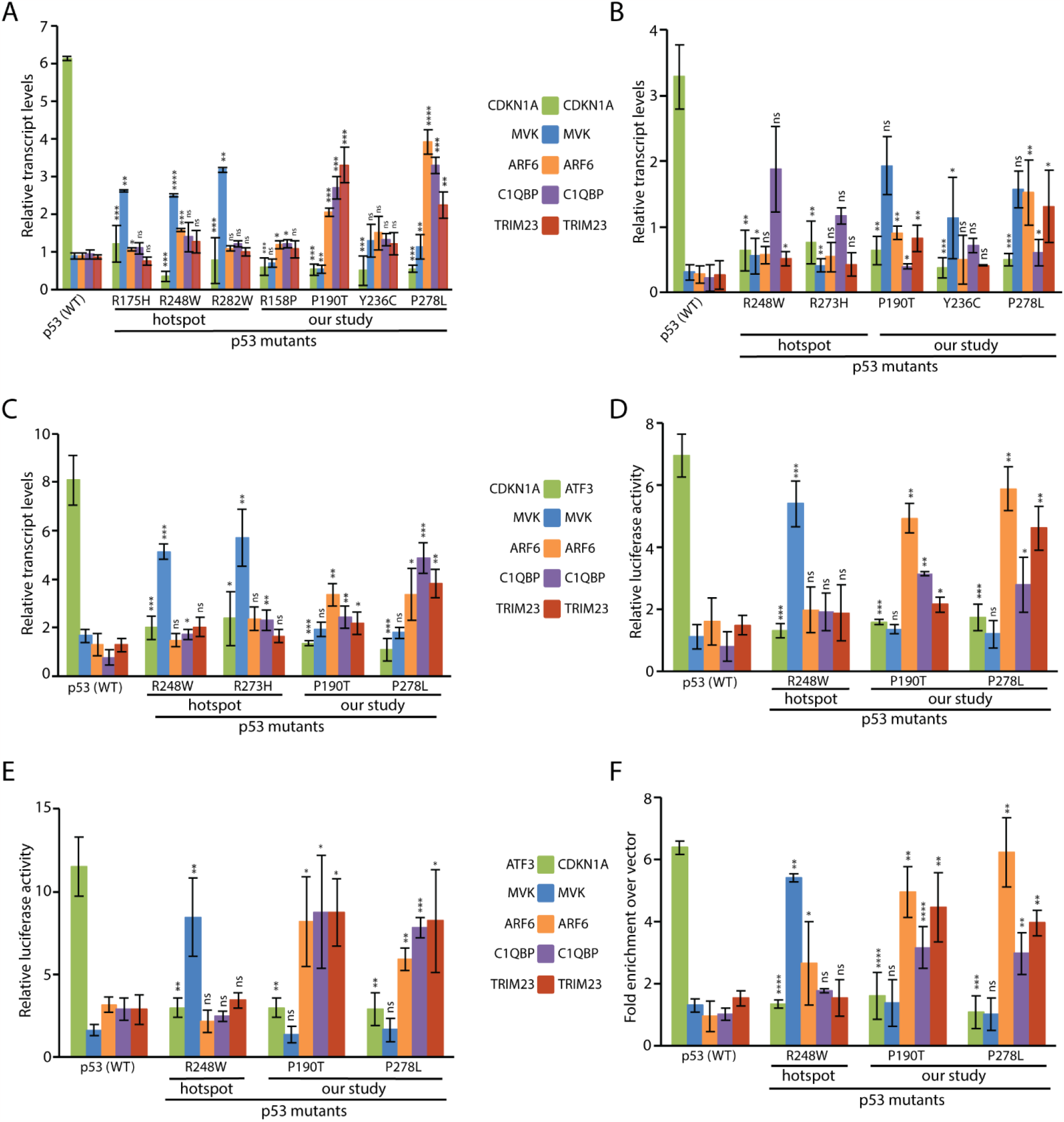
Validation of novel transcriptional targets of specific non-hotspot mutant p53 forms. Results for RT-qPCR assay in H1299 (A), AW13516 (B) and KYSE-410 (C); promoter-luciferase assay in H1299 (D) and KYSE-410 (E); ChAP assay in H1299 (F). Each result is based on at least three independent experiments; *, P<0.05; **, P<0.01; ***, P<0.001; ****, P<0.0001; ns, not significant (unpaired student’s t-test).

### The transcriptional targets of non-hotspot mutant p53 play oncogenic roles in squamous cell carcinoma cell lines

Protein over-expression studies in cell lines revealed that P190T and P278L induced the expression of the three selected target genes. Given the ability of mutant p53 to induce oncogenic function in tumor cells via transcriptional activation of relevant genes, we proceeded to determine if *ARF6, C1QBP* and *TRIM23* could exhibit pro-oncogenic functions in squamous cancer cell lines. To this end, we performed shRNA-based knockdown studies in the ESCC cell line KYSE-410 (Figures 4 and S6) as well as in AW13516 and H1299 cells (Figures S6 and S7). Knockdown of *ARF6, C1QBP* and *TRIM23* resulted in a reduction in several tumorigenic properties assessed by evaluating cell viability, cell growth, colony formation and migratory potential (Figures 4 and S7). Over-expression of the respective proteins resulted in a significant rescue of the original tumorigenic properties of the cell lines, thus confirming the specificity of results obtained with knockdown of each of the three genes (Figures 5, S6, S8 and S9). Interestingly, ectopic expression in control cells (generated from scramble shRNA and thereby not exhibiting any knockdown) also revealed significant activation of tumorigenic properties further confirming the potential oncogenic roles of *ARF6, C1QBP* and *TRIM23* (Figures 5, S6, S8 and S9).

**Figure 4.**
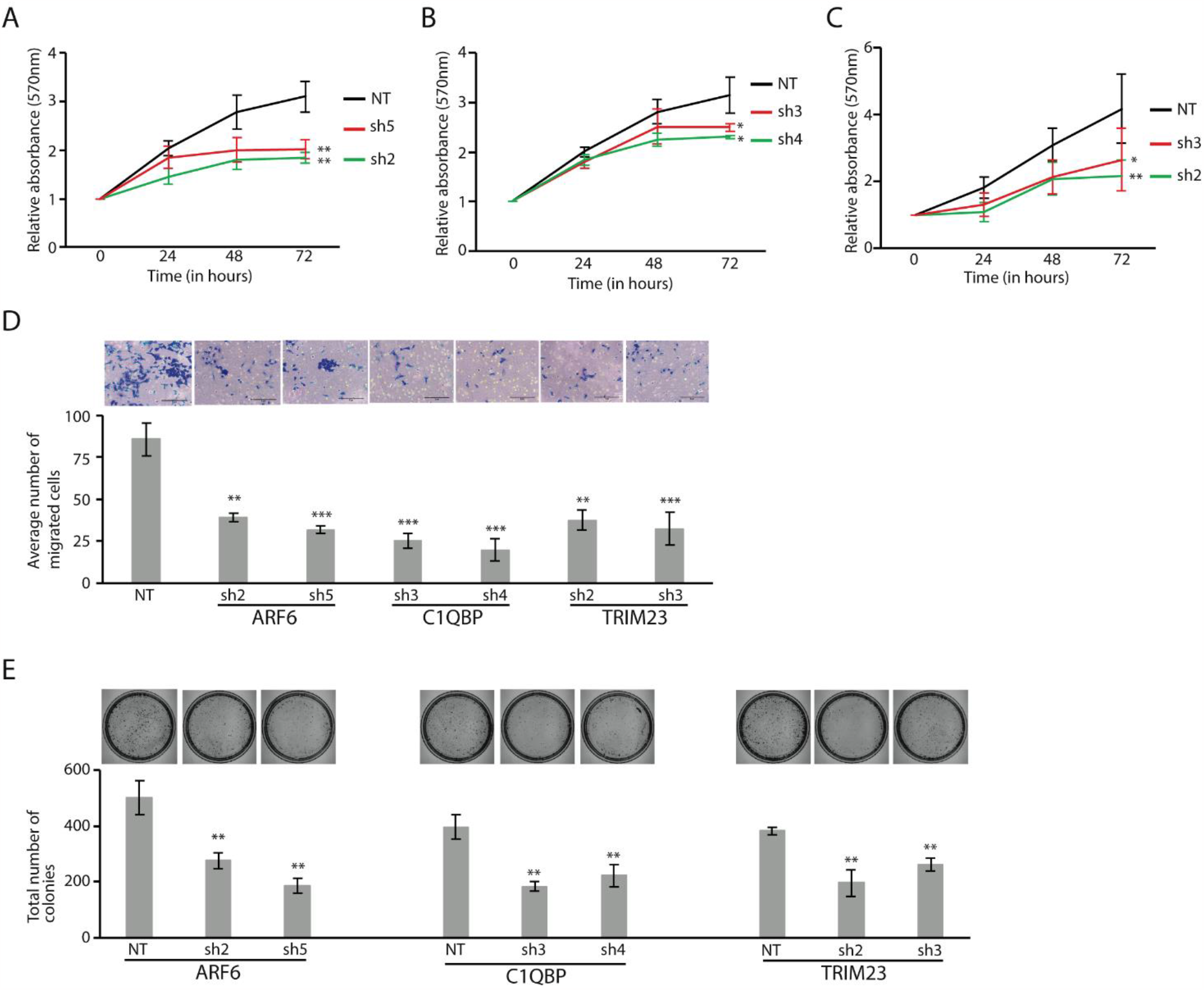
*ARF6, C1QBP, TRIM23* exhibit oncogenic properties in KYSE-410 cells as determined by performing various tumorigenic assays following stable knockdown. Results for various tumorigenic assays following stable knockdown of each of the three genes is shown. MTT (A-C); cell migration (D); colony formation (E). Each result is based on at least three independent experiments; *, P<0.05; **, P<0.01; ***, P<0.001 (unpaired student’s t-test).

**Figure 5.**
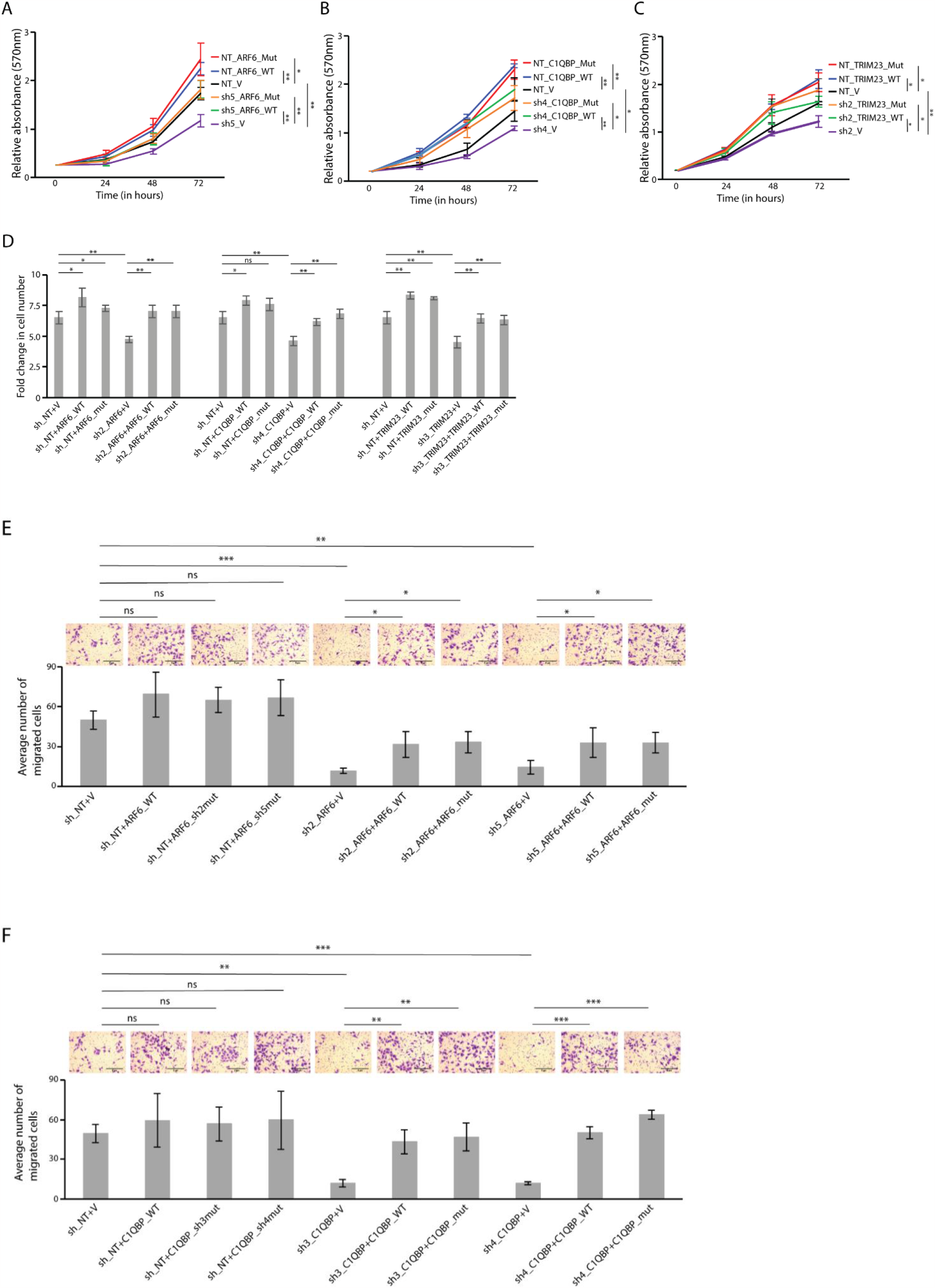
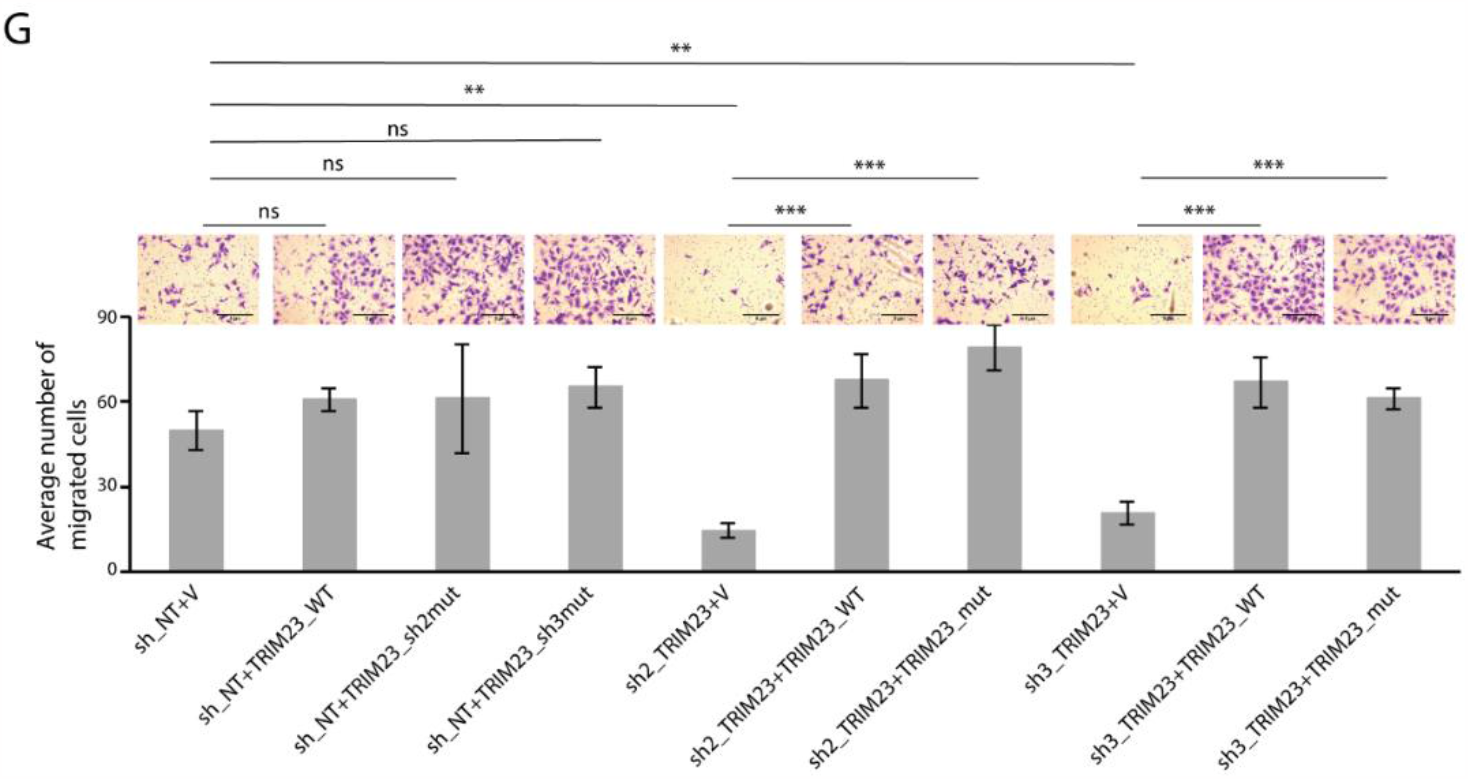
Rescue of reduced cell viability (A-C), cell growth (D) and migration (E-F) upon respective protein expression in KYSE-410 cells harboring stable knockdown of *ARF6, C1QBP* or *TRIM23*. Note, rescue is obtained upon ectopic expression of both wild-type and mutant (generated to negate effect of shRNA) proteins. Each result is based on at least three independent experiments; *, P<0.05; **, P<0.01; ***, P<0.001; ns, not significant (unpaired student’s t-test).

Next, we evaluated the oncogenic potential of *C1QBP* and *TRIM23* in nude mice xenograft models. AW13516 cells exhibiting knockdown of each gene resulted in significant reduction in tumor size and volume in comparison to the control cells (Figures 6 and S10).

**Figure 6.**
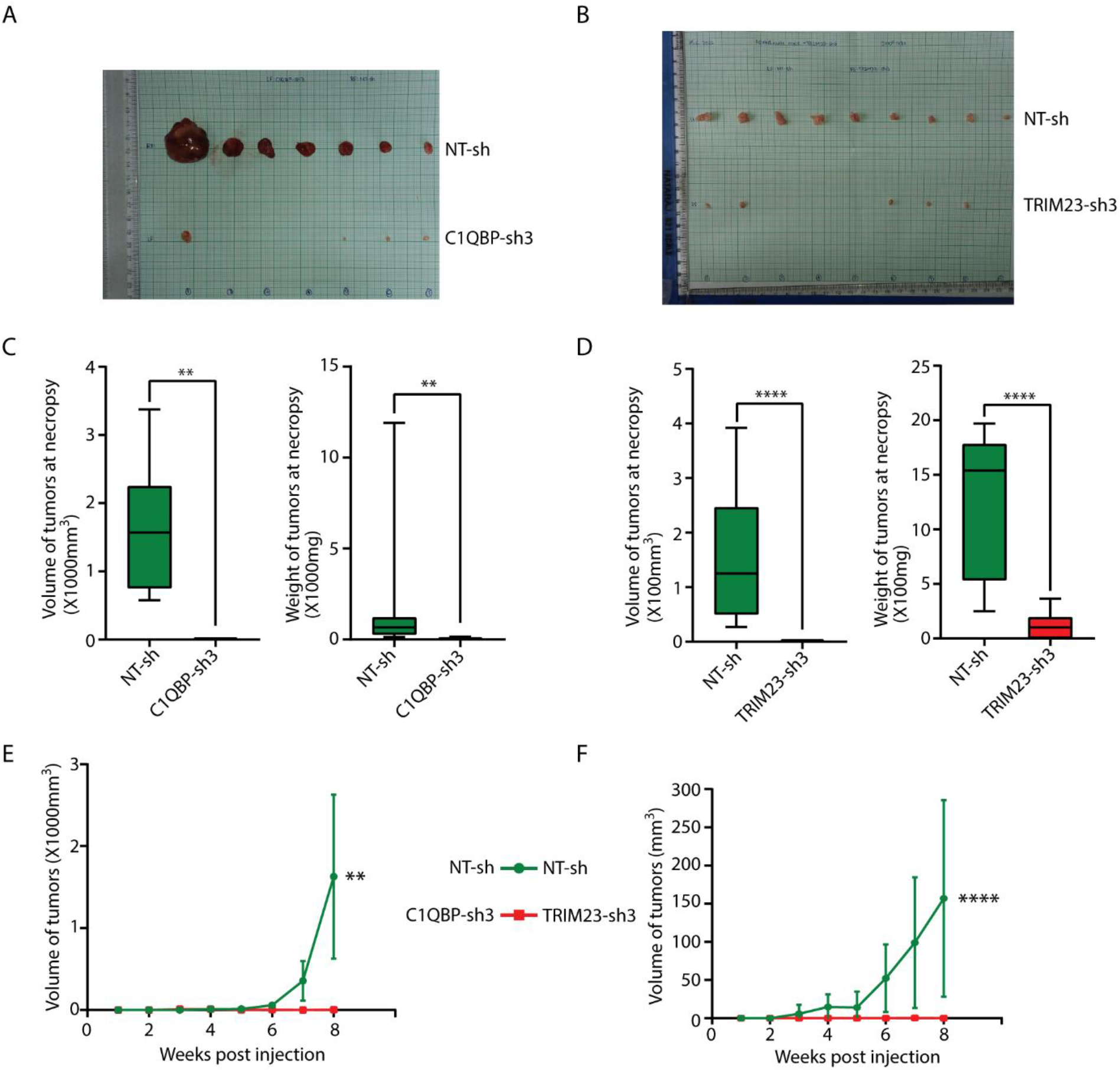
Validation of the oncogenic role of C1QBP (A, C and E) and TRIM23 (B, D and F) in AW13516 squamous carcinoma cells based on nude mice xenograft assays. Representative images of tumors excised from the animals (A-B) and quantitation of their volume and weight (C-D) are shown. The periodic change in tumor volume is shown in panels E-F. **, P<0.01; ****, P<0.0001 (unpaired student’s t-test).

We assessed *ARF6, C1QBP* and *TRIM23* transcript levels in ESCC tumor compared to normal samples. As expected, the transcript levels of all three genes were significantly higher in the tumors in comparison to the matched normal samples, further validating the potential oncogenic nature of these genes (Figure 7A; Table S2D).

**Figure 7.**
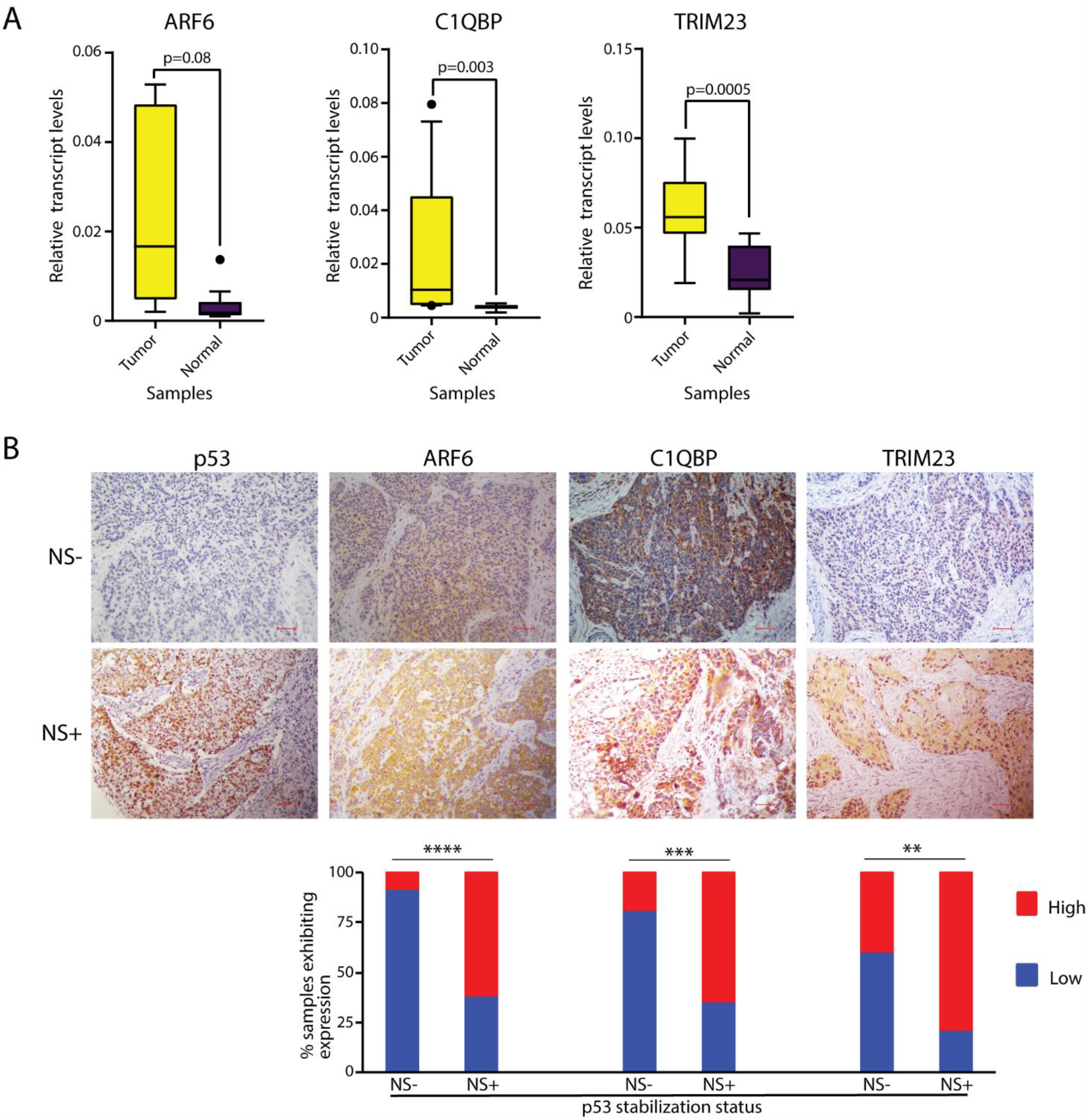
Validation of the oncogenic role of ARF6, C1QBP and TRIM23 from patient-derived tumor samples. Quantitation of relative transcript levels of the three genes in tumor vs. normal ESCC samples (A). Correlation of expression of the three genes with that of p53 using IHC on an ESCC tissue microarray (B).

Finally, the protein expression and localization of p53 and its target gene products was determined by immunohistochemistry on ESCC tumor samples (Figure 7B). A positive correlation was observed between the nuclear stabilization of p53 and the expression of the mutant p53 transcriptional target proteins – ARF6, C1QBP and TRIM23 (Figure 7B and Table S4). These findings suggest a clinical relevance of the discovery of novel transcriptional targets of mutant p53.

## Discussion

Several recent studies have revealed the oncogenic potential of p53 mutant forms that frequently occur (‘hotspot’ mutants) in various cancers. The oncogenic potential exhibited by hotspot p53 mutant forms such as R175H, R273H and R248Q/W has been shown to result from their ability to transcriptionally activate genes with the ability to induce oncogenic changes in cells (Muller & Vousden, 2014). In contrast, evaluation of the potential oncogenic roles of less-frequent non-hotspot p53 mutants has been lacking. The main objective of the current study was therefore to evaluate the oncogenic potential of these rare p53 mutant forms identified from ESCC tumors and also to characterize the hitherto uncharacterized transcriptional networks they may regulate.

The ability to induce oncogenic changes in cells does not appear to be a common feature of all mutant p53 forms, rather only specific mutants seem to exhibit this ability (Dittmer et al., 1993; Hsiao et al., 1994). Indeed, in our study, R158P and Y236C exhibited weak or no oncogenic activity when compared to hotspot mutants whereas P278L and P190T exhibited robust oncogenic functions.

Based on a comprehensive analysis, we identified three potential oncogenic transcriptional targets (*ARF6, C1QBP* and *TRIM23*) of rare p53 mutant forms (P190T and P278L) with high relevance to ESCC. The ARF6 protein controls cell adhesion, actin remodelling, cytokinesis and membrane trafficking (D’Souza-Schorey & Chavrier, 2006; Donaldson, 2003; Grossmann et al., 2019). High levels of *ARF6* and its downstream targets have been reported in cancers of the breast, pancreas, liver and lung, and have been linked to elevated invasive, proliferative and metastatic potential of the tumors (Hashimoto et al., 2004; Liang et al., 2017; Tsutaho et al., 2020). ARF6 and its effector protein AMAP1 form the EGFR-GEP100-ARF6-AMAP1 axis to induce invasion in various cancers (Hashimoto et al., 2016; Sato et al., 2014) besides disturbing the cell adhesion regulated by E-cadherin (Xu et al., 2015). C1QBP levels are elevated in cancers of the breast, prostate, thyroid, esophagus, endometrium, pancreas, stomach, colon and lung (Li et al., 2017; Matsumoto & Bay, 2021). C1QBP levels are found to exhibit a strong correlation with metastasis and high tumor stage in breast cancer, and are associated with higher stage and recurrence in prostate cancer (Amamoto et al., 2011; X. Zhang et al., 2013). TRIM23 is an E3 ubiquitin ligase that localizes to lysosomes and golgi vesicles and plays a role in vesicular transport and phospholipase-D activation (Meroni & Diez-Roux, 2005). *TRIM23* transcript levels were found to be high in cancers of the stomach, liver, lung and colorectum (Han et al., 2020; Y. Zhang et al., 2020). TRIM23 activates NF-κB in hepatocellular carcinoma cells leading to the suppression of HNF-1α and in turn of miR-194 resulting in increased invasion and migration (Bao et al., 2015).

Therefore, in this study, we identified P278L and P190T as novel gain of function non-hotspot p53 mutations presumably involved in ESCC tumorigenesis. More importantly, three novel and exclusive transcriptional targets namely *ARF6, C1QBP* and *TRIM23* of P278L and P190T were identified which exhibit pro-oncogenic properties indicating their direct or indirect role in the development of squamous cell carcinomas. It is essential to define the complete transcriptional network of different p53 mutants which is expected to aid in the development of suitable therapeutic regimes to efficiently treat cancer patients. We believe that the work done in this study is an important step in that direction.

## Acknowledgements

The authors thank all the patients who consented to the study. The authors are grateful to Dr Mukta Srinivasulu of the Mehdi Nawaz Jung Institute of Oncology & Regional Cancer Centre, Hyderabad; Drs Mohana Vamsy Chigurupati and Snehalatha Dhagam of Omega Hospitals, Hyderabad; Drs Sachin S Marda and Srihari of Yashoda Hospitals, Hyderabad; Dr Satish Rao of Krishna Institute of Medical Sciences, Hyderabad and Dr Umanath K Nayak of the Apollo Hospitals, Hyderabad for facilitating sample collection, providing clinical data and assessing molecular data. We acknowledge Dr Raju SR Adduri, former PhD student in the Laboratory of Molecular Oncology, CDFD, Hyderabad for assisting with the computational analysis of the microarray data. We thank Ms. Sanjana Sarkar for the cloning of shRNA and generation of viral particles for transduction experiments. The authors also thank Ms Leena Bashyam and Prof Aparna DuttaGupta, Genomics Facility, University of Hyderabad, Hyderabad, for scanning and initial data extraction of microarrays. We also thank Drs Amit Dutt and Abhijit De from the Advanced Centre for the Treatment, Research and Education in Cancer, Mumbai, India; Dr Rinu Sharma from the Guru Gobind Singh Indraprasta University, New Delhi, India; Dr Saumitra Das from the Indian Institute of Science, Bengaluru, India; Dr Rashna Bhandari from the Centre for DNA Fingerprinting and Diagnostics, Hyderabad, India; and Drs Akhilesh Pandey and Aditi Chatterjee, Institute of Bioinformatics, Bengaluru, India for kindly providing us with cell lines.

## Author contributions

MDB conceived, arranged funding and supervised the study. MDB, SAG and PR designed the experiments. SAG, VK and PR performed experiments. SG and SGU evaluated and scored the H&E and immunohistochemistry data. SAG, VK, PR and MDB analysed and interpreted the results. MDB and SAG wrote the manuscript

