## Supplementary data for "Identification of novel oncogenic transcriptional targets of mutant p53 in Esophageal Squamous Cell Carcinoma"

**Supplementary files:**

**Table S1. Details of samples used for array-based expression analysis of ESCC samples**

| Sample number | Age/ Gender | Grade | Stage | IHC status | p53 status |
|---|---|---|---|---|---|
| 3215 | 22/M | I | pT3N2 | NS- | WT |
| 3244 | 45/M | I | pT3N0 | NS- | WT |
| 3251 | 52/F | I | pT3N1 | NS- | WT |
| 3259 | 50/F | I | pT3N1 | NS- | WT |
| 3270 | 34/F | I | pT3N1 | NS- | WT |
| 3272 | 55/M | I | pT3N2 | NS- | WT |
| 3313 | 48/F | II | pT3N1bM1b | NS- | WT |
| 3325 | 64/M | II | pT2N1a | NS- | WT |
| 3351 | 70/M | II | pT3N1 | NS- | WT |
| 3353 | 68/F | II | pT2N0Mx | NS+ | p.G266R |
| 3357 | 23/F | II | pT2N2 | NS- | WT |
| 3361 | 45/M | II | NA | NS- | WT |
| 3367 | 62/M | I | pT3N0 | NS+ | WT |
| 3369 | 59/M | I | pT2N0 | NS+ | p.P278L |
| 3375 | 64/F | II | pT2N1b | NS- | WT |
| 3379 | 58/F | II | pT3N1Mx | NS+ | WT |
| 3385 | 50/F | II | pT3N1 | NS- | WT |
| 3389 | 41/M | I | pT3N1 | NS+ | p.F134C |
| 3391 | 50/M | I | pT3N2Mx | NS- | WT |
| 3395 | 47/M | I | pT2N1 | NS+ | WT |
| 3401 | 66/F | I | pT3N2Mx | NS+ | WT |
| 3417 | 62/M | II | pT3N0Mx | NS+ | p.R158P |
| 3421 | 38/F | I | pT3N1 | NS+ | p.P190T |
| 3441 | 79/M | I | pT2N0 | NS- | WT |
| 3453 | 61/M | I | pT3N1 | NS- | WT |
| 3467 | 38/M | I | pT3N0 | NS+ | p.P152T |
| 3479 | 59/M | I | pT3N0 | NS+ | p.C176Y |
| 3485 | 39/F | I | pT2N0 | NS+ | p.V172F |
| 3503 | 32/F | I | pT3N2Mx | NS- | WT |

| | | | | | |
|---|---|---|---|---|---|
| 3505 | 55/F | I | pT3N1 | NS- | WT |
| 3511 | 46/M | I | pT3N1 | NS+ | WT |
| 3519 | 69/M | II | pT3N0 | NS+ | p.R248W* |
| 3521 | 61/M | I | pT2N1 | NS+ | WT |
| 3529 | 54/M | II | pT2N1 | NS+ | p.Y163C |
| 3547 | 38/M | I | pT3N0 | NS+ | p.Y236C |
| 3561 | 64/M | I | pT3N0 | NS- | WT |

*, a hotspot mutation

**Table S2A.** Number of samples used for RT-qPCR based gene expression validation in p53 NS+ vs NS- samples.

| **Gene** | **Number of samples** | | | | | |
|---|---|---|---|---|---|---|
| | **Arrayed** | | **Non-arrayed** | | **Total** | |
| | **p53 status** | | **p53 status** | | **p53 status** | |
| | **NS+** | **NS-** | **NS+** | **NS-** | **NS+** | **NS-** |
| *ARF6* | 8 | 10 | 2 | 3 | 10 | 13 |
| *C1QBP* | 6 | 9 | 4 | 2 | 10 | 11 |
| *TRIM23* | 7 | 10 | 3 | 3 | 10 | 13 |

**Table S2B.** Number of samples used for RT-qPCR based transcript level correlation between *TP53* and *ARF6*, *C1QBP* and *TRIM23*.

| **Gene** | **Number of samples** | | |
|---|---|---|---|
| | **Arrayed** | **Non-arrayed** | **Total** |
| *TP53* | 22 | 5 | 27 |
| *ARF6* | 22 | 5 | 27 |
| *C1QBP* | 16 | 3 | 19 |
| *TRIM23* | 22 | 5 | 27 |

**Table S2C.** Number of samples used for RT-qPCR based gene expression validation in p53 mutant vs wild type samples.

| **Gene** | **Number of samples** | | | | | |
|---|---|---|---|---|---|---|
| | **Arrayed** | | **Non-arrayed** | | **Total** | |
| | **p53 status** | | **p53 status** | | **p53 status** | |
| | **WT** | **Mutant** | **WT** | **Mutant** | **WT** | **Mutant** |
| *ARF6* | 11 | 7 | 5 | 0 | 16 | 7 |
| *C1QBP* | 9 | 6 | 4 | 2 | 13 | 8 |
| *TRIM23* | 11 | 6 | 6 | 0 | 17 | 6 |

**Table S2D.** Number of samples used for RT-qPCR based gene expression validation in ESCC tumor vs matched normal.

| **Gene** | **Number of samples** | | | | | |
|---|---|---|---|---|---|---|
| | **Arrayed** | | **Non-arrayed** | | **Total** | |
| | **p53 status** | | **p53 status** | | **p53 status** | |
| | **Tumour** | **Normal** | **Tumour** | **Normal** | **Tumour** | **Normal** |
| *ARF6* | 8 | 8 | 4 | 4 | 12 | 12 |
| *C1QBP* | 8 | 8 | 4 | 4 | 12 | 12 |
| *TRIM23* | 8 | 8 | 2 | 2 | 10 | 10 |

**Table S3: Frequency of *TP53* mutations evaluated in this study in ESCC mutation databases.**

| Mutation | The *TP53* Database* | COSMIC# | TCGA~ |
|---|---|---|---|
| Our study | | | |
| F134 | 0.07% (0%) | 0.15% (0%) | 1.04% (0%) |
| P152 | 0.16% (0%) | 0.28% (0%) | 0% (0%) |
| R158 | 0.10% (0.03%) | 0.35% (0.03%) | 2.08% (0%) |
| Y163 | 0.16% (0.16%) | 0.43% (0.35%) | 1.04% (1.04%) |
| V172 | 0.16% (0.10%) | 0.18% (0.13%) | 0% (0%) |
| C176 | 2.07% (0.10%) | 0.96% (0.30%) | 1.04% (1.04%) |
| P190 | 0.19% (0.07%) | 0.28% (0.01%) | 0% (0%) |
| Y236 | 0.26% (0.23%) | 0.46% (0.38%) | 0% (0%) |
| R248† | 1.87% (0.94%) | 1.82% (0.01%) | 3.13% (0%) |
| G266 | 0.45% (0.19%) | 0.35% (0.20%) | 0% (0%) |
| P278 | 0.65% (0.13%) | 0.68% (0.15%) | 4.17% (0%) |
| Hotspot mutants | | | |
| R175 | 1.91% | 2.00% | 6.25% |
| G245 | 1.16% | 1.39% | 1.04% |
| R249 | 0.32% | 0.56% | 2.08% |
| R273 | 1.71% | 1.49% | 6.25% |
| R282 | 1.26% | 1.54% | 3.13% |

* https://dceg.cancer.gov/tools/public-data/tp53-database

### https://cancer.sanger.ac.uk/cosmic

~ https://gdac.broadinstitute.org/runs/stddata__2016_01_28/data/ESCA/20160128/

† a hotspot mutant

The values in brackets indicate the percentage of the mutants identified in our study, ie. F134C, P152T, R158P, Y163C, V172F, C176Y, P190T, Y236C, R248W, G266R and P278L, in the respective databases.

**Table S4. IHC-based evaluation of expression of mutant p53 target genes in ESCC samples.**

| Sample | IHC result | |
|---|---|---|
| | p53 status[a] | p53 target protein (staining score) |
| ARF6[b] | | |
| 3215 | NS- | 2 |
| 3244 | NS- | 1 |
| 3246 | NS- | 1 |
| 3251 | NS- | 1 |
| 3259 | NS- | 2 |
| 3265 | NS- | 2 |
| 3270 | NS- | 1 |
| 3272 | NS- | 1 |
| 3276 | NS- | 1 |
| 3280 | NS- | 1 |
| 3293 | NS- | 1 |
| 3301 | NS- | 1 |
| 3315 | NS- | 1 |
| 3321 | NS- | 4 |
| 3351 | NS- | 2 |
| 3361 | NS- | 2 |
| 3379 | NS- | 1 |
| 3391 | NS- | 1 |
| 3441 | NS- | 4 |
| 3445 | NS- | 1 |
| 3447 | NS- | 2 |
| 3457 | NS- | 1 |
| 3491 | NS- | 1 |
| 3493 | NS- | 1 |
| 3495 | NS- | 1 |
| 3501 | NS- | 2 |
| 3503 | NS- | 1 |
| 3505 | NS- | 1 |

| | | |
|---|---|---|
| 3507 | NS- | 1 |
| 3511 | NS- | 1 |
| 3521 | NS- | 1 |
| 3525 | NS- | 1 |
| 3561 | NS- | 2 |
| 3587 | NS- | 6 |
| 3595 | NS- | 1 |
| 3249 | NS+ | 6 |
| 3274 | NS+ | 2 |
| 3278 | NS+ | 6 |
| 3282 | NS+ | 4 |
| 3286 | NS+ | 1 |
| 3298 | NS+ | 2 |
| 3313 | NS+ | 1 |
| 3317 | NS+ | 4 |
| 3323 | NS+ | 4 |
| 3325 | NS+ | 4 |
| 3345 | NS+ | 6 |
| 3347 | NS+ | 6 |
| 3349 | NS+ | 4 |
| 3353 | NS+ | 4 |
| 3367 | NS+ | 1 |
| 3369 | NS+ | 4 |
| 3385 | NS+ | 2 |
| 3387 | NS+ | 6 |
| 3389 | NS+ | 6 |
| 3395 | NS+ | 4 |
| 3397 | NS+ | 4 |
| 3399 | NS+ | 6 |
| 3401 | NS+ | 2 |
| 3403 | NS+ | 2 |
| 3407 | NS+ | 4 |
| 3413 | NS+ | 4 |
| 3415 | NS+ | 4 |

| | | |
|---|---|---|
| 3417 | NS+ | 1 |
| 3421 | NS+ | 4 |
| 3427 | NS+ | 4 |
| 3449 | NS+ | 1 |
| 3451 | NS+ | 6 |
| 3467 | NS+ | 2 |
| 3479 | NS+ | 4 |
| 3487 | NS+ | 4 |
| 3519 | NS+ | 1 |
| 3529 | NS+ | 2 |
| 3549 | NS+ | 2 |
| C1QBP[c] | | |
| 3215 | NS- | 5 |
| 3246 | NS- | 1 |
| 3251 | NS- | 2 |
| 3259 | NS- | 5 |
| 3265 | NS- | 4 |
| 3270 | NS- | 1 |
| 3272 | NS- | 2 |
| 3276 | NS- | 1 |
| 3280 | NS- | 2 |
| 3301 | NS- | 2 |
| 3321 | NS- | 0 |
| 3351 | NS- | 2 |
| 3361 | NS- | 2 |
| 3379 | NS- | 1 |
| 3391 | NS- | 4 |
| 3441 | NS- | 1 |
| 3445 | NS- | 3 |
| 3447 | NS- | 2 |
| 3457 | NS- | 1 |
| 3491 | NS- | 2 |
| 3493 | NS- | 2 |

| | | |
|---|---|---|
| 3495 | NS- | 1 |
| 3501 | NS- | 2 |
| 3503 | NS- | 1 |
| 3505 | NS- | 2 |
| 3507 | NS- | 2 |
| 3511 | NS- | 1 |
| 3521 | NS- | 2 |
| 3525 | NS- | 1 |
| 3561 | NS- | 5 |
| 3587 | NS- | 2 |
| 3595 | NS- | 2 |
| 3249 | NS+ | 3 |
| 3274 | NS+ | 3 |
| 3278 | NS+ | 4 |
| 3286 | NS+ | 3 |
| 3298 | NS+ | 2 |
| 3313 | NS+ | 5 |
| 3317 | NS+ | 4 |
| 3323 | NS+ | 2 |
| 3325 | NS+ | 2 |
| 3345 | NS+ | 4 |
| 3347 | NS+ | 4 |
| 3353 | NS+ | 5 |
| 3367 | NS+ | 3 |
| 3369 | NS+ | 1 |
| 3385 | NS+ | 3 |
| 3387 | NS+ | 3 |
| 3389 | NS+ | 3 |
| 3395 | NS+ | 3 |
| 3397 | NS+ | 5 |
| 3399 | NS+ | 3 |
| 3401 | NS+ | 1 |
| 3403 | NS+ | 2 |
| 3407 | NS+ | 4 |

| | | |
|---|---|---|
| 3415 | NS+ | 4 |
| 3417 | NS+ | 1 |
| 3427 | NS+ | 2 |
| 3449 | NS+ | 3 |
| 3451 | NS+ | 3 |
| 3467 | NS+ | 4 |
| 3479 | NS+ | 3 |
| 3529 | NS+ | 1 |
| TRIM23[d] | | |
| 215 | NS- | 1 |
| 244 | NS- | 1 |
| 246 | NS- | 1 |
| 251 | NS- | 5 |
| 259 | NS- | 7 |
| 265 | NS- | 7 |
| 270 | NS- | 1 |
| 272 | NS- | 1 |
| 276 | NS- | 1 |
| 280 | NS- | 6 |
| 3301 | NS- | 6 |
| 3321 | NS- | 6 |
| 3351 | NS- | 2 |
| 3361 | NS- | 3 |
| 3379 | NS- | 3 |
| 3391 | NS- | 6 |
| 3441 | NS- | 6 |
| 3445 | NS- | 1 |
| 3447 | NS- | 1 |
| 3457 | NS- | 5 |
| 3491 | NS- | 3 |
| 3493 | NS- | 1 |
| 3495 | NS- | 5 |
| 3501 | NS- | 2 |

| | | |
|---|---|---|
| 3503 | NS- | 1 |
| 3505 | NS- | 1 |
| 3507 | NS- | 1 |
| 3511 | NS- | 6 |
| 3521 | NS- | 6 |
| 3525 | NS- | 1 |
| 3561 | NS- | 1 |
| 3587 | NS- | 1 |
| 3595 | NS- | 5 |
| 249 | NS+ | 6 |
| 274 | NS+ | 4 |
| 278 | NS+ | 6 |
| 286 | NS+ | 6 |
| 298 | NS+ | 7 |
| 3313 | NS+ | 4 |
| 3317 | NS+ | 7 |
| 3323 | NS+ | 7 |
| 3325 | NS+ | 4 |
| 3345 | NS+ | 6 |
| 3347 | NS+ | 1 |
| 3349 | NS+ | 2 |
| 3353 | NS+ | 4 |
| 3367 | NS+ | 1 |
| 3369 | NS+ | 1 |
| 3385 | NS+ | 6 |
| 3387 | NS+ | 6 |
| 3389 | NS+ | 6 |
| 3395 | NS+ | 1 |
| 3397 | NS+ | 6 |
| 3399 | NS+ | 6 |
| 3401 | NS+ | 3 |
| 3403 | NS+ | 5 |
| 3407 | NS+ | 6 |
| 3413 | NS+ | 6 |

| | | |
|---|---|---|
| 3415 | NS+ | 6 |
| 3417 | NS+ | 2 |
| 3421 | NS+ | 6 |
| 3427 | NS+ | 6 |
| 3449 | NS+ | 3 |
| 3451 | NS+ | 6 |
| 3467 | NS+ | 6 |
| 3479 | NS+ | 6 |
| 3487 | NS+ | 7 |
| 3519 | NS+ | 6 |
| 3529 | NS+ | 6 |

[a], NS-, nuclear unstable p53; NS+, nuclear-stabilized p53

[b], Score: 0-3, low expression; 4-6, high expression

[c], Score: 0-2, low expression; 3-5, high expression

[d], Score: 0-3, low expression; 4-7, high expression

A

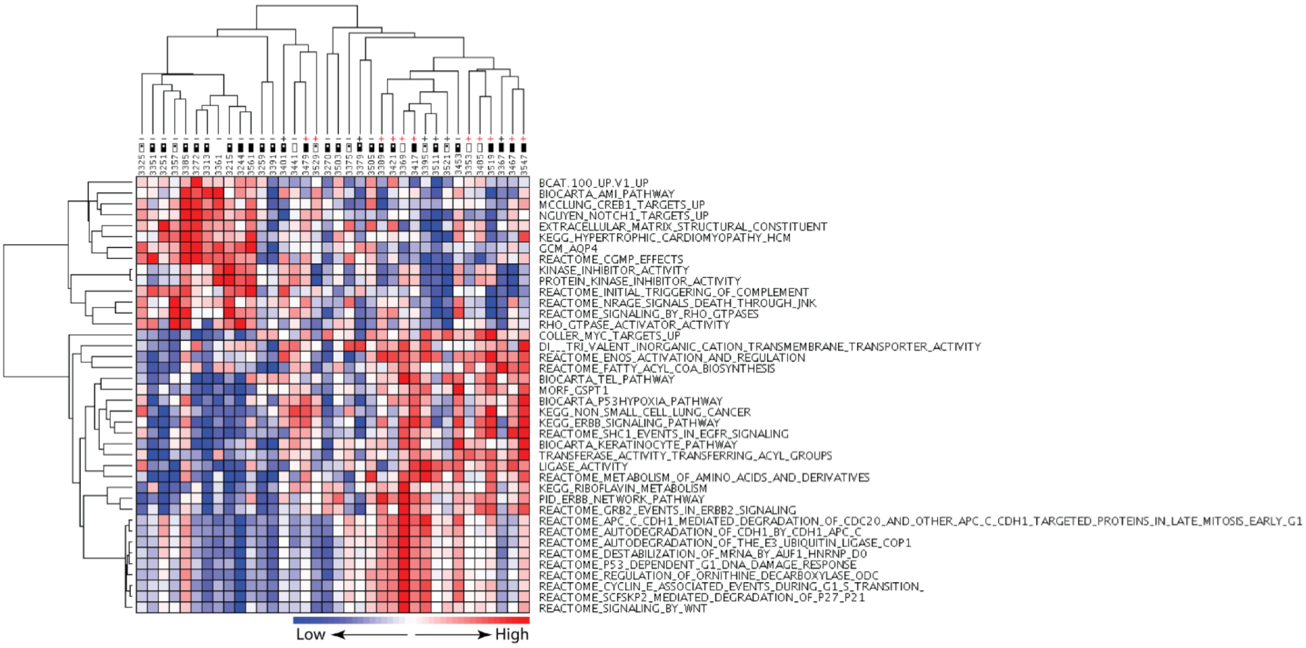


B

| Expression line | Name | Description | Expected score | Observed score | Numerator | Denominator | Fold change | q-value (%) |
|---|---|---|---|---|---|---|---|---|
| **Genes upregulated in tumors exhibiting p53 nuclear stabilization** | | | | | | | | |
| | ARF6 | na | -1.4865668 | 3.6902077 | 2.0453758 | 0.55427116 | 2.5122864 | 0.0 |
| | C1QBP | na | -1.0394305 | 3.65616 | 1.3123622 | 0.3589455 | 2.7961118 | 0.0 |
| | WASL | na | 2.2478516 | 3.6343613 | 1.3567052 | 0.37329948 | 1.8622425 | 0.0 |
| | TRIM23 | na | 1.7440538 | 3.880726 | 3.1665344 | 0.8159644 | 2.5837252 | 0.0 |
| | GOT2 | na | -0.21365492 | 4.5545573 | 1.9896443 | 0.4368469 | 4.3247514 | 0.0 |
| | TP53 | na | 1.6352164 | 4.16442 | 1.7121012 | 0.41112596 | 6.483293 | 0.0 |
| | VDAC1 | na | 2.219359 | 3.9761484 | 2.272691 | 0.57158107 | 2.0142248 | 0.0 |
| **Genes upregulated in tumors not exhibiting p53 nuclear stabilization** | | | | | | | | |
| | DLC1 | na | -0.55462176 | -4.4632187 | -2.0073254 | 0.4497484 | 0.123302184 | 0.0 |
| | TPM1 | na | 1.653539 | -4.286875 | -2.2658107 | 0.52854604 | 0.18994986 | 0.0 |
| | CACNG8 | na | -0.9526078 | -4.2228622 | -2.1591077 | 0.51129013 | 0.11761566 | 0.0 |
| | NR2F1 | na | 0.30004066 | -3.8630028 | -2.3682604 | 0.613062 | 0.053999852 | 0.0 |
| | SGCD | na | 0.9742601 | -3.7810917 | -2.547929 | 0.67386067 | 0.20978683 | 0.0 |
| | ITGA9 | na | -0.016006574 | -3.725075 | -2.4248638 | 0.65095705 | 0.19806127 | 0.0 |
| | RHOBTB1 | na | 0.82734925 | -3.630808 | -2.928195 | 0.8064858 | 0.33126834 | 0.0 |
| | EFEMP1 | na | -0.503197 | -3.6007752 | -3.0186563 | 0.8383351 | 0.3318192 | 0.0 |
| | LAMC1 | na | 0.062020887 | -3.4968803 | -1.8726773 | 0.535528 | 0.2415704 | 0.0 |
| | ANXA6 | na | -1.5231924 | -3.493382 | -1.3649838 | 0.3907342 | 0.33641902 | 0.0 |

C

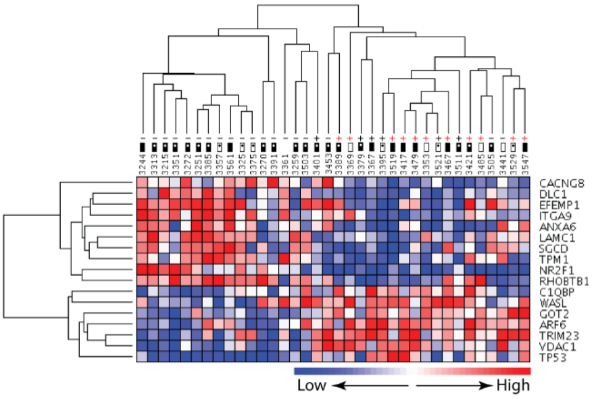


**Figure S1. Genome-wide transcript profiling of ESCC tumor samples to identify transcriptional targets of mutant p53.** Identification of gene sets (A) and genes (B-C) differentially expressed between ESCC tumor samples stratified for p53 status. – and +, p53 NS- and NS+ respectively (red colour denotes mutant p53); box with and without dot, node positive and negative respectively; open and filled box, tumor stage 1 or 2 and 3 or 4, respectively.

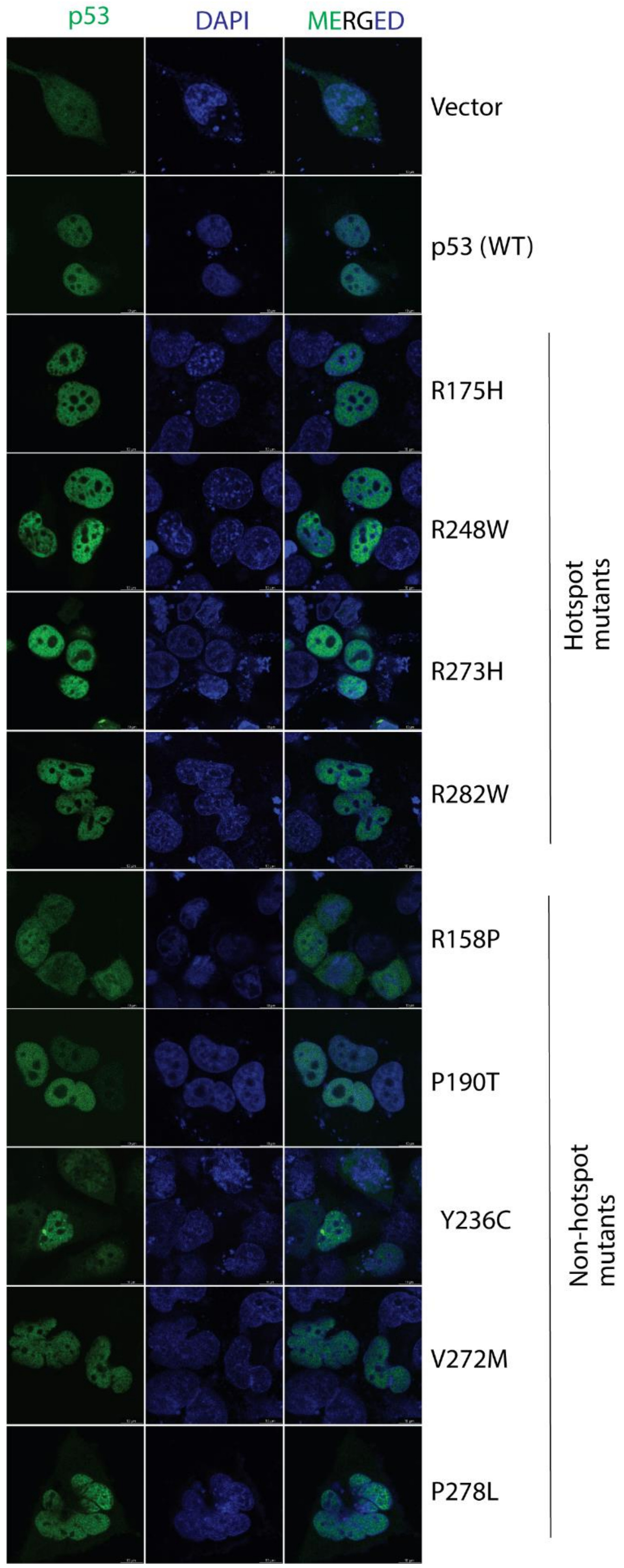


**Figure S2. Fluorescence-based visualization of intracellular localization of wild type and various mutant p53 forms.** The localizations of different ectopically-expressed p53-EGFP proteins were assessed in H1299 squamous cell carcinoma cell line.

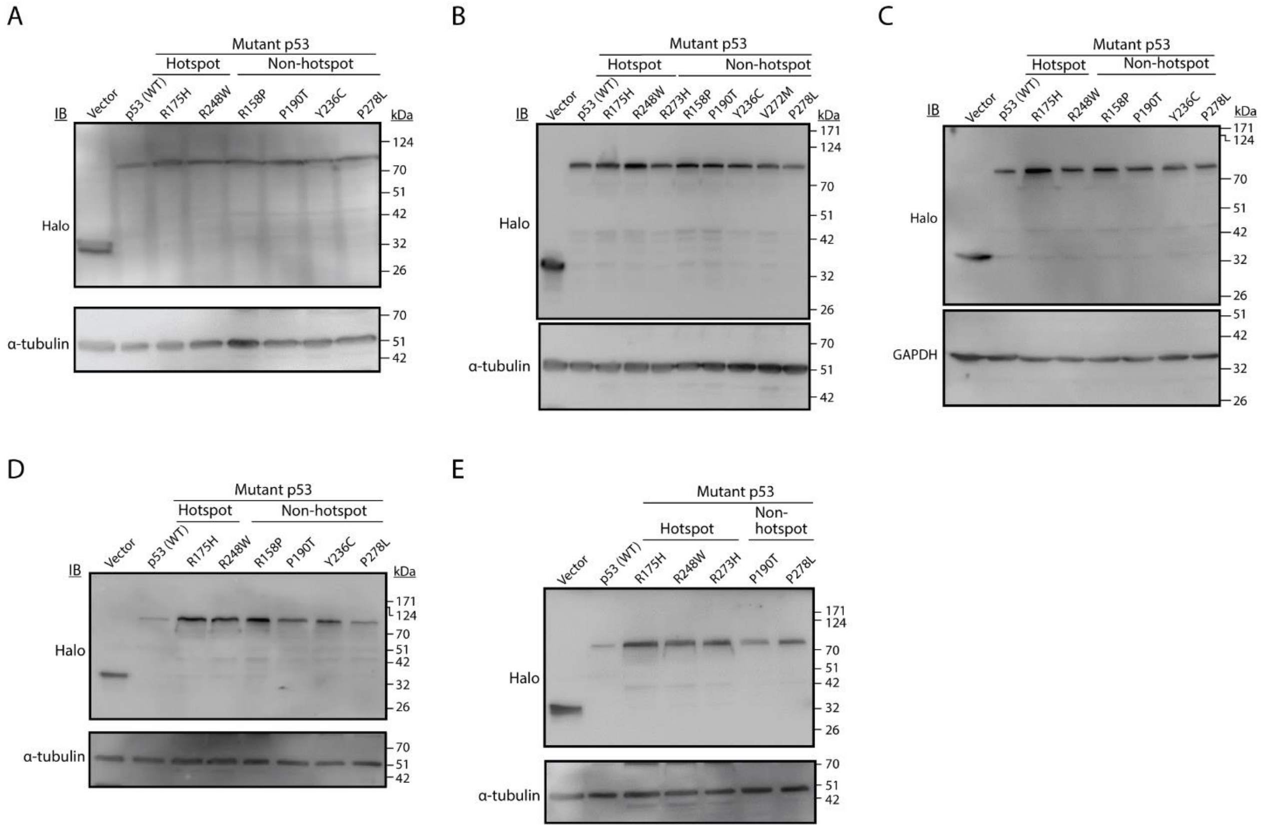


**Figure S3. Evaluation of expression levels of ectopically expressed wild type and different mutant p53 forms in KYSE-410 (A), H1299 (B), NT8e (C), AW13516 (D) and JHU-011 (E) cells; in relation to Figures 2, 3 and S4.**

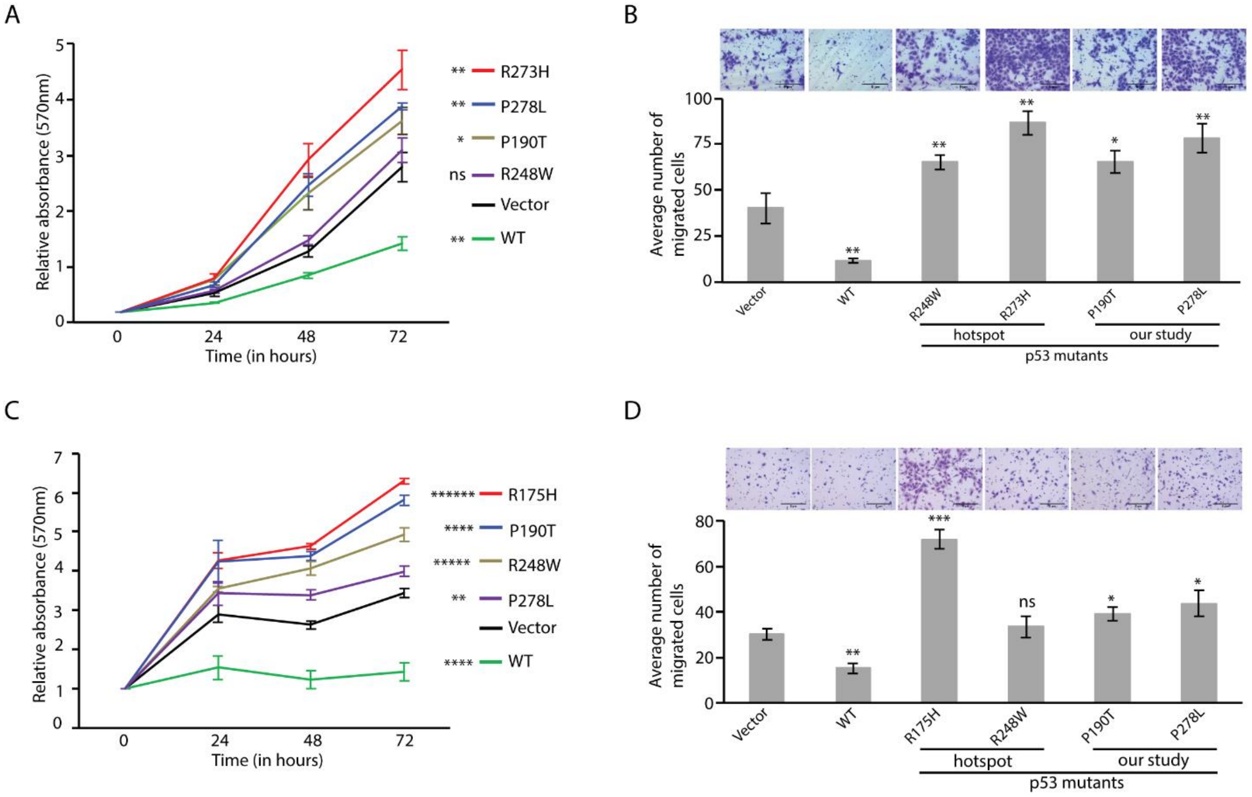


**Figure S4. Evaluation of oncogenic potential of 'non-hotspot' mutant p53 proteins (in relation to Figure 2). Results for MTT (A, C) and cell migration (B, D) are shown for NT8e (A-B) and JHU-011 (C-D) cells.** Each result is based on at least three independent experiments; *, $P<0.05$; **, $P<0.01$; ***, $P<0.001$; ****, $P<0.0001$; *****, $P<0.00001$; ******, $p<0.000001$; ns, not significant (unpaired student's t-test).

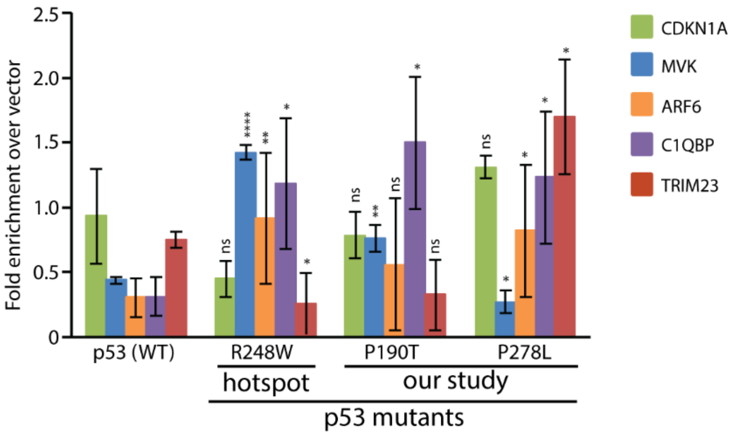


**Figure S5. ChAP assay for gene body region of target genes for wild type and various mutant p53 forms (in relation to Figure 3F).** Each result is based on at least three independent experiments; *, P<0.05; **, P<0.01; ****, P<0.0001; ns, not significant (unpaired student's t-test).

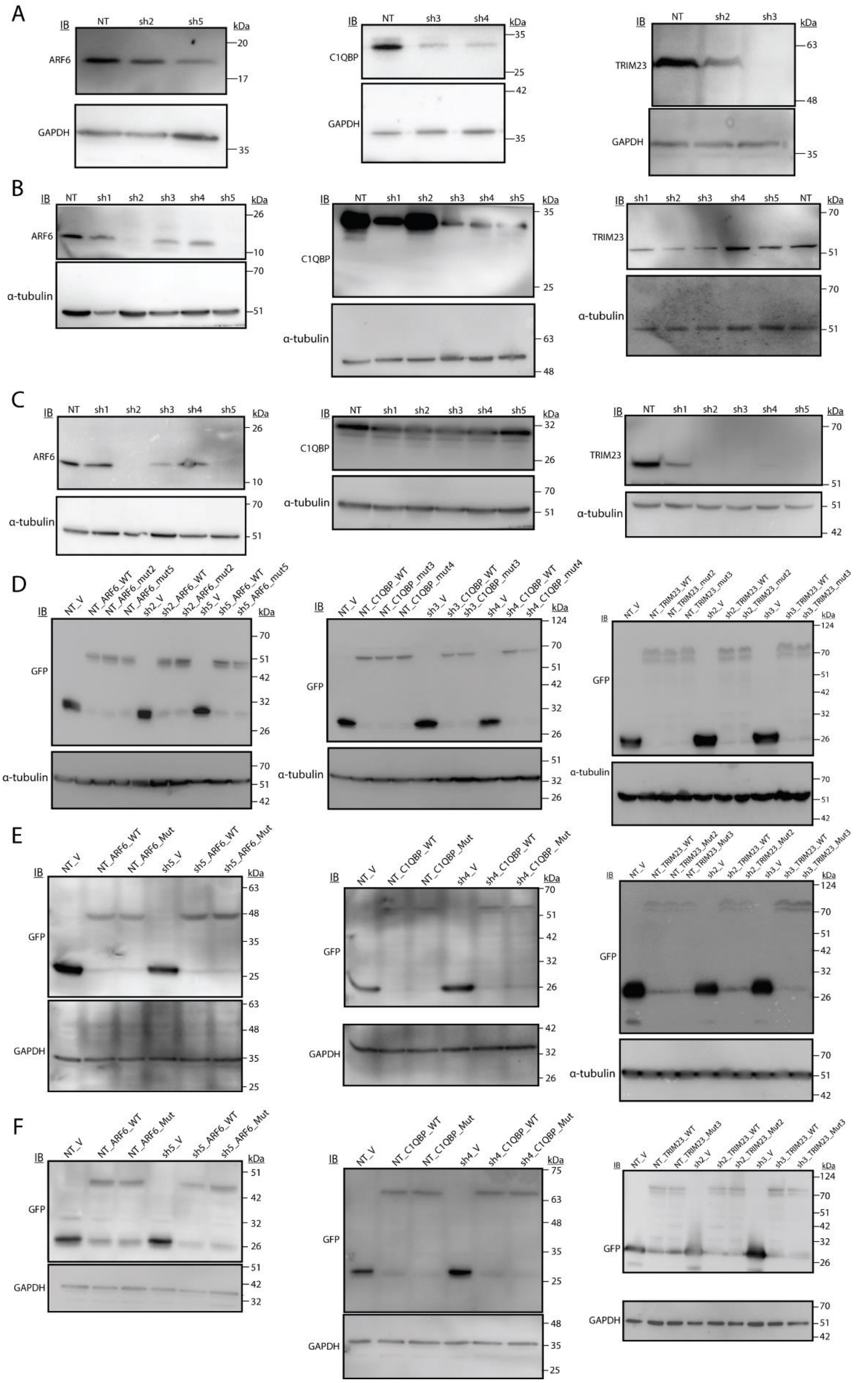


**Figure S6. Evaluation of the expression levels of ARF6, C1QBP or TRIM23 to confirm knockdown (A-C; in relation to Figure 4 and S7), or expression in rescue experiments (D-F) in relation to Figures 5, S8 and S9.** A, D, KYSE-410; B, E, AW13516; C, F, H1299.

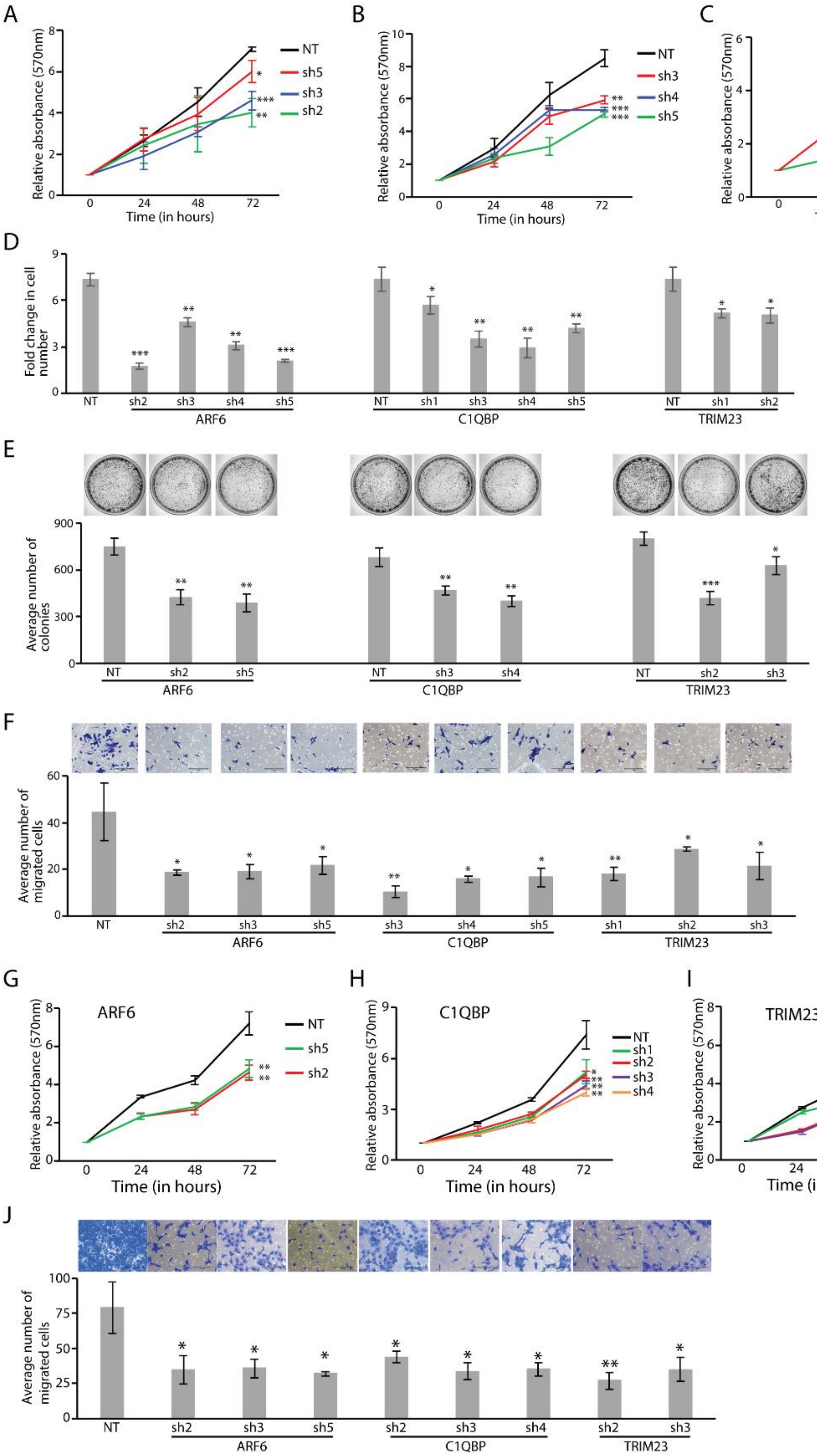
A
Relative absorbance (570nm)
Time (in hours)
NT
sh5
sh3
sh2
B
Relative absorbance (570nm)
Time (in hours)
NT
sh3
sh4
sh5
C
Relative absorbance (570nm)
Time (in hours)
NT
sh2
sh1
D
Fold change in cell number
NT
sh2
sh3
sh4
sh5
ARF6
NT
sh1
sh3
sh4
sh5
C1QBP
NT
sh1
sh2
TRIM23
E
Average number of colonies
NT
sh2
sh5
ARF6
NT
sh3
sh4
C1QBP
NT
sh2
sh3
TRIM23
F
Average number of migrated cells
NT
sh2
sh3
sh5
ARF6
sh3
sh4
sh5
C1QBP
sh1
sh2
sh3
TRIM23
G
ARF6
Relative absorbance (570nm)
Time (in hours)
NT
sh5
sh2
H
C1QBP
Relative absorbance (570nm)
Time (in hours)
NT
sh1
sh2
sh3
sh4
I
TRIM23
Relative absorbance (570nm)
Time (in hours)
NT
sh4
sh2
sh3
J
Average number of migrated cells
NT
sh2
sh3
sh5
ARF6
sh2
sh3
sh4
C1QBP
sh2
sh3
TRIM23

**Figure S7. ARF6, C1QBP and TRIM23 exhibit oncogenic properties in AW13516 (A-F) and H1299 (G-I) cells as determined by performing various tumorigenic assays following stable knockdown (in relation to Figure 4).** MTT (A-C and G-I); cell growth (D); colony formation (E), and cell migration (F and J). Each result is based on at least three independent experiments; *, $P<0.05$; **, $P<0.01$; ***, $P<0.001$; ns, not significant (unpaired student's t-test).

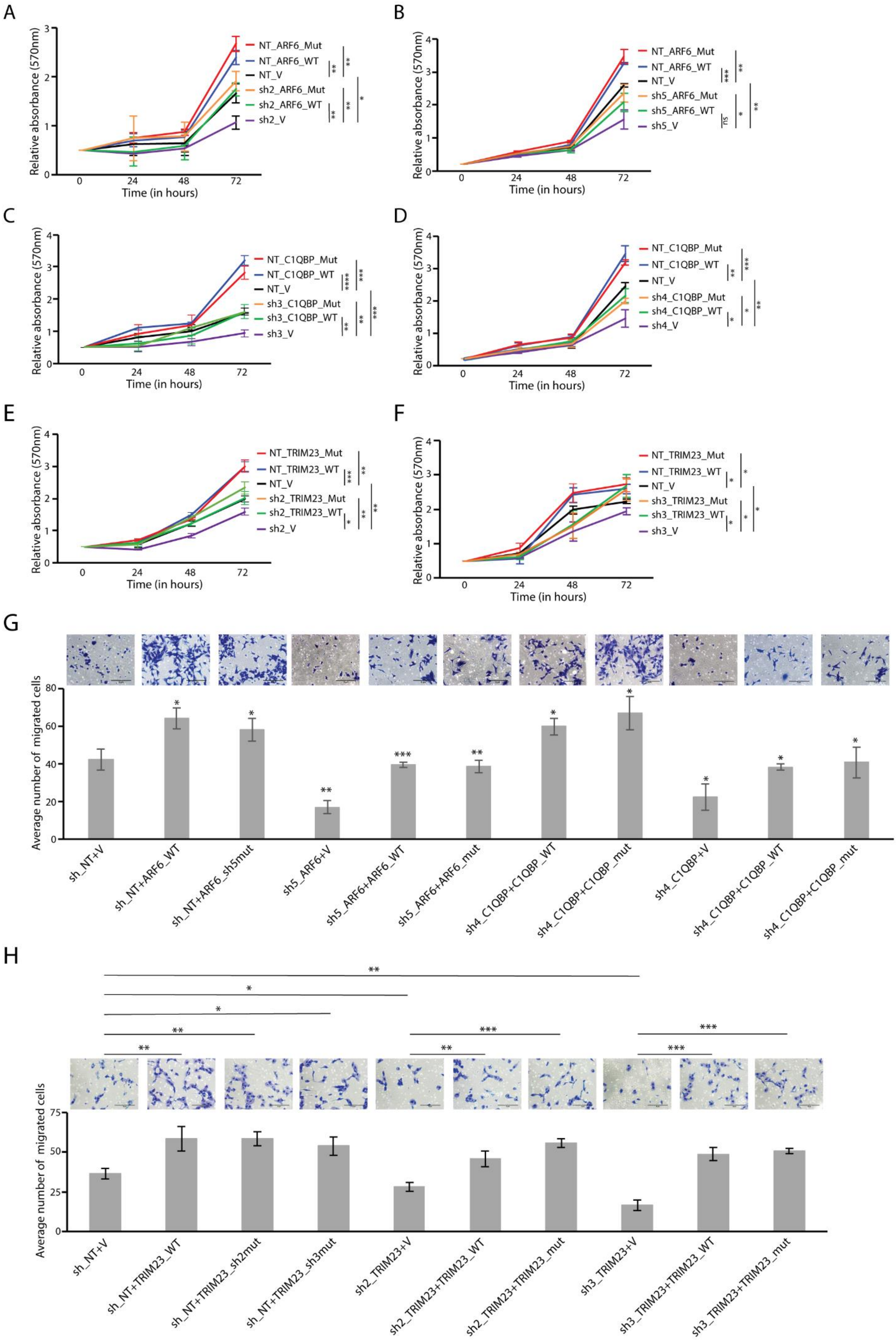
A
Relative absorbance (570nm)
Time (in hours)
NT_ARF6_Mut
NT_ARF6_WT
NT_V
sh2_ARF6_Mut
sh2_ARF6_WT
sh2_V
B
NT_ARF6_Mut
NT_ARF6_WT
NT_V
sh5_ARF6_Mut
sh5_ARF6_WT
sh5_V
ns
C
NT_C1QBP_Mut
NT_C1QBP_WT
NT_V
sh3_C1QBP_Mut
sh3_C1QBP_WT
sh3_V
D
NT_C1QBP_Mut
NT_C1QBP_WT
NT_V
sh4_C1QBP_Mut
sh4_C1QBP_WT
sh4_V
E
NT_TRIM23_Mut
NT_TRIM23_WT
NT_V
sh2_TRIM23_Mut
sh2_TRIM23_WT
sh2_V
F
NT_TRIM23_Mut
NT_TRIM23_WT
NT_V
sh3_TRIM23_Mut
sh3_TRIM23_WT
sh3_V
G
Average number of migrated cells
sh_NT+V
sh_NT+ARF6_WT
sh_NT+ARF6_sh5mut
sh5_ARF6+V
sh5_ARF6+ARF6_WT
sh5_ARF6+ARF6_mut
sh4_C1QBP+C1QBP_WT
sh4_C1QBP+C1QBP_mut
sh4_C1QBP+V
sh4_C1QBP+C1QBP_WT
sh4_C1QBP+C1QBP_mut
H
Average number of migrated cells
sh_NT+V
sh_NT+TRIM23_WT
sh_NT+TRIM23_sh2mut
sh_NT+TRIM23_sh3mut
sh2_TRIM23+V
sh2_TRIM23+TRIM23_WT
sh2_TRIM23+TRIM23_mut
sh3_TRIM23+V
sh3_TRIM23+TRIM23_WT
sh3_TRIM23+TRIM23_mut

**Figure S8. Rescue of reduced cell viability (A-F) and cell migration (G-H) upon respective protein expression in AW13516 cells harboring stable knockdown of *ARF6, C1QBP* or *TRIM23*.** Both the wild type and the mutant (generated to avoid being targeted by the shRNA) forms of the over-expressed target proteins were observed to rescue the loss of oncogenic phenotypes brought about by their respective knockdown. Each result is based on at least three independent experiments; *, P<0.05; **, P<0.01; ***, P<0.001; ****, P<0.0001; ns, not significant (unpaired student's t-test).

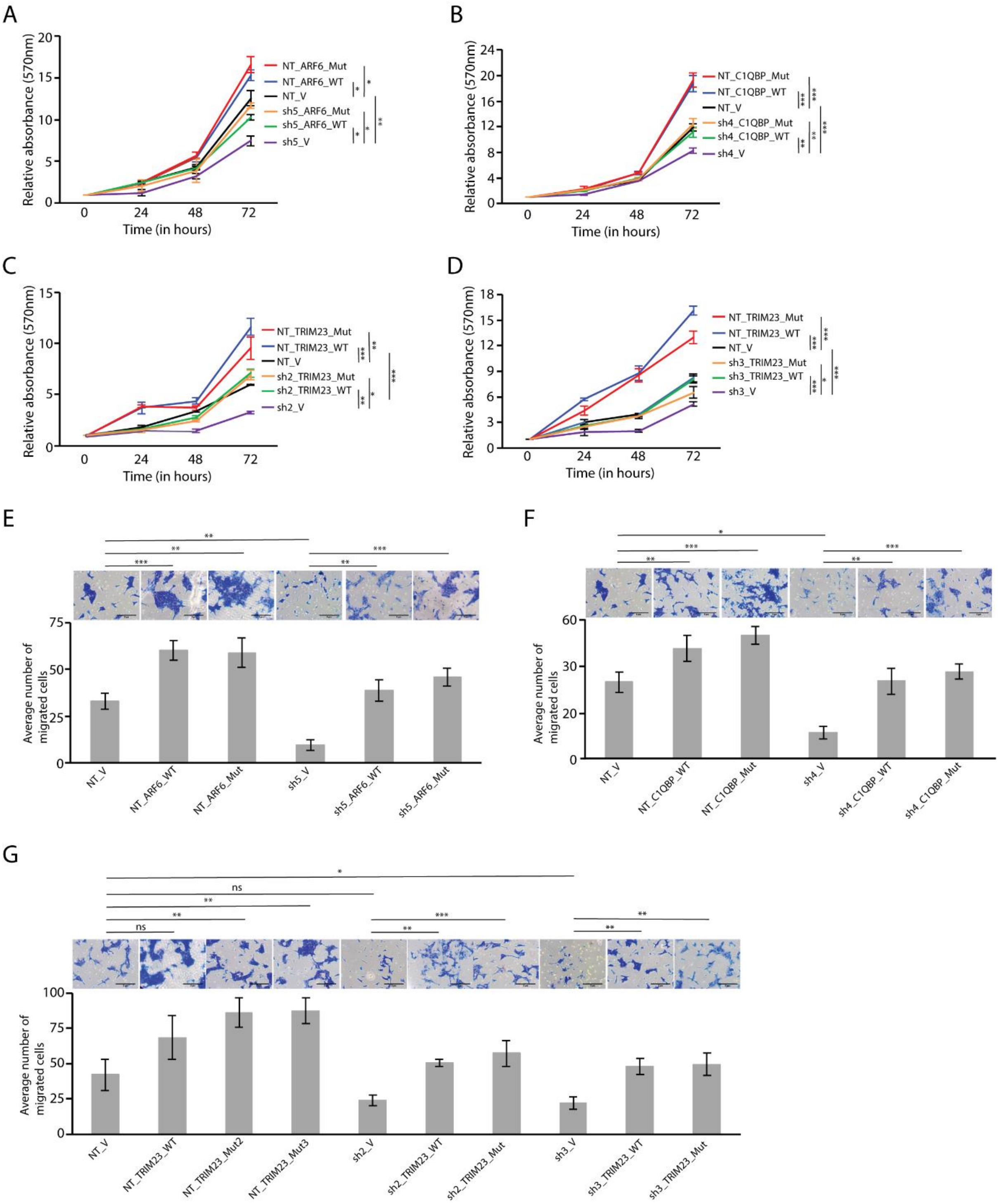


**Figure S9. Rescue of reduced cell viability (A-D) and cell migration (E-G) upon respective protein expression in H1299 cells harboring stable knockdown of *ARF6*, *C1QBP* or *TRIM23*.** Both the wild type and the mutant (generated to avoid being targeted by the shRNA) forms of the over-expressed target proteins were observed to rescue the loss of oncogenic phenotypes brought about by their respective knockdown. Each result is based on at least three independent experiments; *, P<0.05; **, P<0.01; ***, P<0.001; ns, not significant (unpaired student's t-test).

A

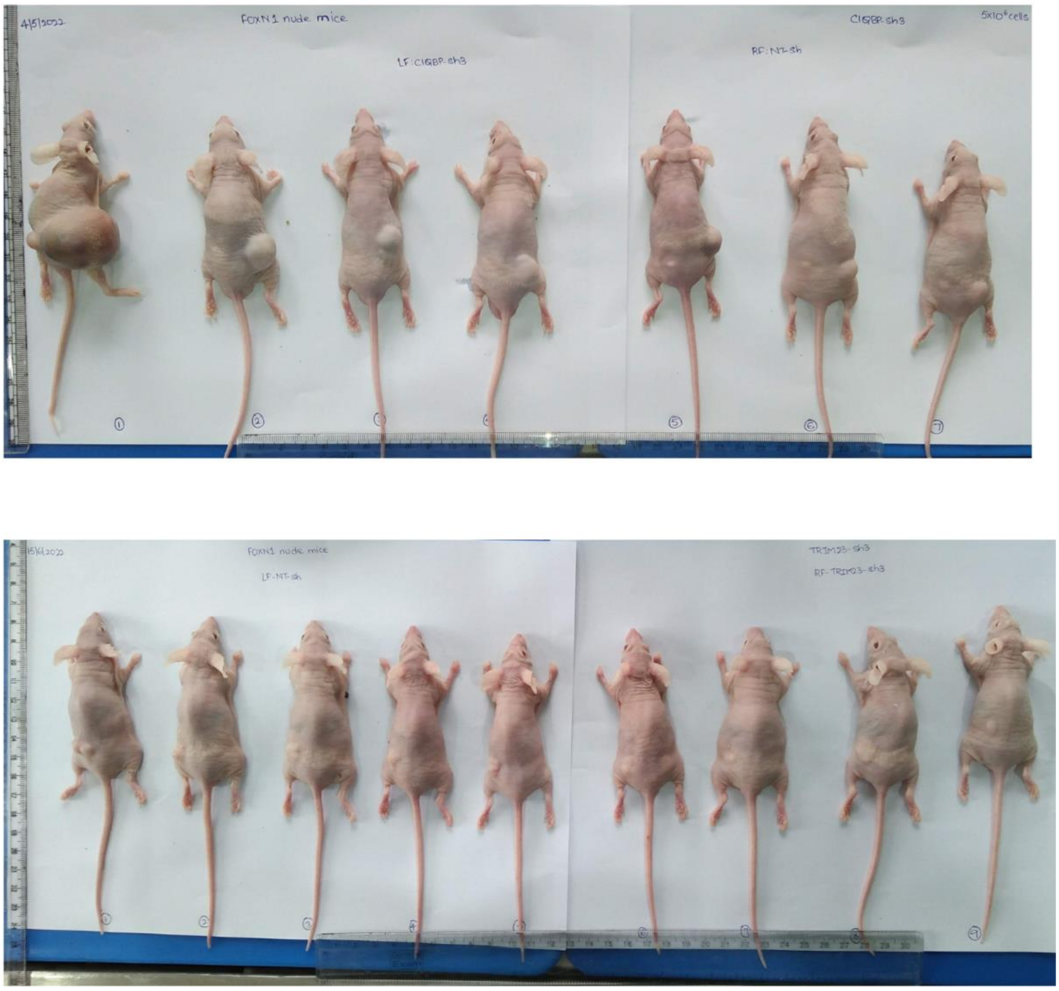

**Figure S10. Validation of the oncogenic role of C1QBP and TRIM23 in AW13516 squamous carcinoma cells based on nude mice xenograft assays**. **(in relation to Figure 6).** Images of all mice injected with AW13516 cells harboring either control (right flank: C1QBP (panel A), left flank: TRIM23 (panel B)) or gene-specific (left flank: C1QBP (panel A), right flank: TRIM23 (panel B)) shRNA stable constructs (C1QBP, panel A; TRIM23, panel B) following euthanization is shown.
